# Select azo compounds post-translationally modulate HTRA1 abundance and activity potentially through interactions at the trimer interface

**DOI:** 10.1101/2025.05.13.651909

**Authors:** John D. Hulleman, Seungje Jeon, Sofia Bali, Sophia M. DiCesare, Ali Abbas, Steffi Daniel, Antonio J. Ortega, Gracen E. Collier, Julian Yang, Archishman Bhattacharyaa, Melissa K. McCoy, Lukasz A. Joachimiak, Bruce A. Posner

**Affiliations:** Department of Ophthalmology and Visual Neurosciences, University of Minnesota, 2001 6th St. SE, Minneapolis, Minnesota, 55455, United States; Department of Ophthalmology, University of Texas Southwestern Medical Center, 5323 Harry Hines Blvd, Dallas, Texas, 75390, United States; Center for Alzheimer’s and Neurodegenerative Diseases, O’Donnell Brain Institute, University of Texas Southwestern Medical Center, 5323 Harry Hines Blvd, Dallas, Texas, 75390, United States; Department of Biochemistry, University of Texas Southwestern Medical Center, 5323 Harry Hines Blvd, Dallas, Texas, 75390, United States

## Abstract

High-temperature requirement protein A1 (HTRA1) is a secreted serine protease with diverse substrates, including extracellular matrix proteins, proteins involved in amyloid deposition, and growth factors. Accordingly, HTRA1 has been implicated in a variety of neurodegenerative diseases including a leading cause of blindness in the elderly, age-related macular degeneration (AMD). In fact, genome wide association studies have identified that the 10q26 locus which contains *HTRA1* confers the strongest genetic risk factor for AMD. A recent study has suggested that AMD-associated risk alleles in HTRA1 correlate with a significant age-related defect in HTRA1 synthesis in the retinal pigmented epithelium (RPE) within the eye, possibly accounting for AMD susceptibility. Thus, we sought to identify small molecule enhancers of HTRA1 transcription and/or protein abundance using an unbiased high-throughput screening approach. To accomplish this goal, we used CRISPR/Sp.Cas9 engineering to introduce an 11 amino acid luminescent peptide tag (HiBiT) onto the C-terminus of HTRA1 in immortalized ARPE-19 cells. Editing was very efficient (∼88%), verified by genomic DNA analysis, short interfering RNA (siRNA), and HiBiT blotting. Nineteen-hundred and twenty compounds from two libraries were screened. An azo compound with reported anti-amyloidogenic and cardioprotective activity, Chicago Sky Blue 6B (CSB), was identified as an enhancer of endogenous HTRA1 secretion (2.0 ± 0.3 fold) and intracellular levels (1.7 ± 0.2 fold). These results were counter-screened using HiBiT complement factor H (CFH) edited ARPE-19 cells, verified using HiBiT blotting, and were not due to *HTRA1* transcriptional changes. Importantly, serine hydrolase activity-based protein profiling (SH-ABPP) demonstrated that CSB does not affect HTRA1’s specific activity. However, interestingly, in follow-up studies, Congo Red, another azo compound structurally similar to CSB, also substantially increased intracellular HTRA1 levels (up to 3.6 ± 0.3 fold) but was found to significantly impair HTRA1 enzymatic reactivity (0.45 ± 0.07 fold). Computational modeling of potential azo dye interaction with HTRA1 suggests that CSB and Congo Red can bind to the non-catalytic face of the trimer interface but with different orientation tolerances and interaction energies. These studies identify select azo dyes as HTRA1 chemical probes which may serve as starting points for future HTRA1-centered small molecule therapeutics.

## INTRODUCTION

The explosion of fast and inexpensive methods to analyze large amounts of DNA (e.g., Next-generation sequencing, NGS) has led to the identification of genetic susceptibility loci, disease-causing mutations, and an overall wealth of genetic data which can be correlated with disease progression. Specifically, with respect to vision research, genome-wide association studies (GWAS) have provided a plethora of gene candidates associated with optic disc morphology^1^, myopia^2^, susceptibility to glaucoma^3^, and initiation/progression of age-related macular degeneration (AMD)^4, 5^. AMD is the leading cause of irreversible blindness in the elderly in industrialized nations. It is estimated to currently affect more than 196 million individuals worldwide, a number that is expected to rise to ∼288 million by 2040^6^.

While it is clear that AMD is a multifactorial disease which is heavily influenced by one’s environment (e.g., cigarette smoking, and Western diet)^7^, single nucleotide polymorphisms (SNPs) in at least 52 genetic loci can significantly influence one’s AMD risk^4, 8^. One particular locus that confers the strongest genetic risk for AMD^8^ is located on chromosome 10q26. This locus contains two genes, age-related maculopathy susceptibility factor 2 (*ARMS2*) and high-temperature requirement protein A1 (*HTRA1*)^8^. Due to the proximity of these genes and because they are in high linkage disequilibrium, it has been difficult to understand how (and which) variants in this locus influence AMD risk^9^. Whereas independent studies have indicated that select SNPs increase HTRA1 expression^10^ and that overexpression of HTRA1 leads to vascular diseases resembling exudative (wet) AMD^11, 12^, polypoidal choroidal vasculopathy (PCV)^13^, or retinopathy of prematurity (ROP)^14^ in mice, a more recent study suggested that risk-associated SNPs near *HTRA1* have a more nuanced effect and are context-specific^15^. This latter study found that *HTRA1* mRNA and protein levels were reduced in the retinal pigmented epithelium (RPE) from aged individuals with the rs362127733 risk variant^15^. As a secreted serine hydrolase, lower levels of HTRA1 (or its lower activity) would be predicted to favor the buildup of extracellular matrix proteins involved in AMD progression including EGF-containing fibulin-like extracellular matrix protein 1 (EFEMP1, fibulin-3^16, 17^), fibulin-5^18, 19^, and hemicentin (fibulin-6^20, 21^).

Based on the findings that overexpressed HTRA1 induces significant retinal vascular changes resembling exudative AMD, groups have sought to identify HTRA1 neutralizing antibodies^22^, antibody-based allosteric traps^23^, peptides^24^, or small molecules^25^ to inhibit HTRA1 activity. Yet, few small molecule enhancers or chemical probes^26^ have been identified that could be used to increase HTRA1 abundance, activity or both. To overcome this deficiency in the field, we sought to develop an easy-to-use and sensitive cell-based assay that reflected endogenous HTRA1 production from RPE cells using CRISPR-based genome editing^27^. During the initial years after the advent of CRISPR genome editing, groups successfully appended full-length reporter proteins (e.g., fluorescent proteins^28, 29^) to multiple endogenous POIs for easy detection. However, more recently, to avoid the potentially confounding effects of such large protein reporter tags, groups have used split reporter systems (either mNeon Green [mNG, a fluorescent protein] or Nano Luciferase [NanoBiT]) to increase the efficiency of homology-directed repair (HDR) and to minimize the size of the CRISPR-inserted DNA^30, 31^. In these split reporter systems, the DNA encoding for the small portion of the complementation system (i.e., mNG11 or HiBiT) is appended to a gene of interest (GOI) by CRISPR, while the large portion of the protein complementation system (i.e., mNG1-10 or LgBiT) is produced endogenously or supplied exogenously during the assay stage. While split fluorescent systems like mNG1-10/mNG11 are useful for subcellular localization or morphology studies, split luciferases provide a sensitive means to measure protein production, secretion, and fate (at the cost of sub-ideal, lower resolution bioluminescent imaging)^30, 32, 33^.

Herein we successfully CRISPR-engineered two RPE cell lines producing HiBiT-tagged versions of HTRA1 and a control line endogenously expressing HiBiT complement factor H (CFH, another gene highly associated with AMD^34–37^). We demonstrate that this general HiBiT-tagging approach is efficient, highly specific, and versatile. Not only does it allow for sensitive detection and near real-time monitoring of these proteins produced from their endogenous locus, but it can also be easily scaled for small molecule or genetic high-throughput screening (HTS). HTS on HTRA1 HiBiT cells identified common azo dyes including Chicago Sky Blue 6B (CSB) and Congo Red (Congo R) as compounds that can post translationally regulate endogenous and over-expressed HTRA1 abundance and activity. Importantly, this azo scaffold (or something similar) may serve as a lead pharmacophore for the development of future HTRA1 chemical probes, or even potentially therapeutics for patients at risk of HTRA1-based AMD^15^. More broadly, we envision that a similar HiBiT tagging approach can be easily adapted for other POIs with strong genetic associations to diverse ocular diseases^38, 39^, thereby serving as an effective discovery approach for vision research and new drug development.

## MATERIALS AND METHODS

### Cell culture

Human retinal pigmented epithelial cells (ARPE-19, ATCC CRL-2302, Manassas, VA) were originally propagated in DMEM/F12 (Corning, Corning, NY) media supplemented with 10% fetal bovine serum (FBS, Omega Scientific, Tarzana, CA) and 1% penicillin/streptomycin/glutamine (PSQ, Gibco, Waltham, MA). This media (referred to as full DMEM/F12) was used to scale cultures prior to performing experiments/screening. Prior to high-throughput screening (HTS), and when noted, confluent cells were cultured in differentiation media^40^ (referred to as ‘Nic’ media) for at least one week consisting of MEM Alpha (Sigma, St. Louis, MO) or DMEM (Life Technologies, Carlsbad, CA), GlutaMAX (Gibco), 1% FBS, 1% PS, 1x non-essential amino acids (NEAA, Gibco), N1 (Sigma) or N2 supplement B (Stem Cell, Vancouver, Canada), taurine (0.25 mg/mL, Sigma), hydrocortisone (20 ng/mL, Sigma), triiodo-thyronin (0.013 ng/mL, Sigma), and nicotinamide (10 mM, Stem Cell).

### HiBiT cell line generation

Low passage ARPE-19 cells were genomically edited to introduce a C-terminal VS-<u>HiBiT</u>-GG-hemagglutinin (HA) sequence (VS<u>VSGWRLFKKIS</u>GGYPYDVPDYA) in the *HTRA1* locus (Sup. Fig. 1). The decision of placement of the tag was based on previous demonstration of a myc-6xHis tag placement without compromising protein function^13^ and acceptable CRISPR-related parameters. The *HTRA1* genomic sequence (+/-30 bp) surrounding the signal sequence cleavage site (at Ala22) is extremely GC-rich (83.33%, 79.5% in exon 1 as a whole), which makes designing appropriate N-terminal gRNAs difficult in this region. Therefore, we targeted HTRA1 at the C-terminus. Conversely, a 2xFLAG-VS-<u>HiBiT</u> sequence (DYKDDDDKDYKDDDDKVS<u>VSGWRLFKKIS</u>) was inserted N-terminally immediately after the signal sequence cleavage site of CFH. Introduction of this sequence was predicted to not affect CFH signal sequence cleavage (Sup. Fig. 2A, B) and, in fact, we have previously demonstrated that placement of a 2xFLAG immediately after a signal sequence cut site forces signal sequence processing^38, 41^. To accomplish editing, a CRISPR/Cas9 ribonucleoprotein (RNP) was generated using Alt-R Sp. Cas9 Nuclease V3 (Integrated DNA Technologies, IDT, Coralville, IA) loaded with a crRNA/tracrRNA duplex (gRNA listed in Table S1). This RNP, combined with a single-stranded oligodeoxynucleotide (ssODN) repair template (Table S1) and electroporation enhancer were introduced into ARPE-19 cells using electroporation (1400V, 20 ms, 2 pulses, Neon Transfection System, Life Technologies). After electroporation, cells were incubated in antibiotic-free media containing Homology-Directed Repair (HDR) Enhancer V2 (1 DM, IDT) for 48 h. Heterogenous cultures were expanded and verified as described below.

### Genomic editing validation

Heterogenous HiBiT edited cells were harvested and their genomic DNA (gDNA) was extracted (GenElute, Sigma). gDNA from edited cells and unedited control cells was amplified by exon-specific primers (Table S2) and run on a 2% agarose gel followed by imaging on an Odyssey Fc (LI-COR, Lincoln, NE). Higher molecular weight bands corresponding to edited gDNA were excised, extracted, and sent for Sanger sequencing (Eurofins Genomics, Huntsville, AL). A full agarose gel image of the HTRA1 HiBiT gDNA amplification is presented as Sup. Fig. 5A.

### Short interfering RNA (siRNA)

siRNA was used to confirm that the HiBiT signal originated from editing of the desired locus. To accomplish this, siRNAs (Silencer Select, Life Technologies) were introduced into ARPE-19 cells expressing HiBiT-tagged POIs via reverse transfection.

Briefly, siRNA (100 nM, Table S3) was complexed with 1.6 DL Dharmafect 4 (Horizon Discovery, Cambridge, UK) in 250 DL OptiMEM (Gibco), vortexed for 15 sec, and incubated at RT for 20 min. This siRNA complex was combined with 250 DL of ARPE-19 cell suspension (at 466,666 cells/mL, final siRNA concentration: 50 nM) and incubated overnight in 24 well plates (Corning). Fresh media was provided the next day, followed by 48-72 h of knockdown, and a final media change. The effects of siRNA knockdown on HiBiT-tagged proteins of interest were assessed by a lytic or extracellular HiBiT assay (Promega, Madison, WI) on conditioned media.

### HiBiT blotting

Each HiBiT cell line was grown to confluence in a 10 cm dish, followed by a media change into full Nic media for 3-4 days, and then another media change into serum-free Nic media for an additional 3-4 days. Media (∼10 mL) was then collected and concentrated to 200 DL using an Amicon Ultra 30,000 MWCO filter (Millipore Sigma, St. Louis, MO). For intracellular protein assessment, cells were trypsinized, neutralized, washed in PBS or HBSS and then lysed in RIPA buffer (Santa Cruz, Dallas, TX) supplemented with Halt Protease Inhibitor Cocktail (Pierce, Rockford, IL) and benzonase (MilliporeSigma). Intracellular protein was normalized using a BCA Assay (Pierce). Twenty microliters of concentrated media or an equal amount of intracellular protein was separated on a reducing 4-20% Tris Gly SDS-PAGE gel (140V, 80 min, Life Technologies) and transferred to a nitrocellulose membrane (P0 program, iBlot2, Life Technologies). Blots were washed in Tris Buffered Saline with 0.1% Tween-20 (TBS-T) for at least 30 min (at most overnight) to expose the HiBiT tag. Blots were probed with HiBiT blocking buffer supplemented with LgBiT (1:200) and NanoGlo substrate (1:500) for 5 min at RT, then imaged on an Odyssey Fc (LI-COR) using an appropriate exposure time (30 sec – 10 min). Beta actin (1:10,000, LI-COR) was used as a loading control for intracellular protein and blots were imaged on a LI-COR Odyssey CLx using a near-infrared secondary antibody (LI-COR). Full blot images of the HTRA1 HiBiT blot and beta actin are presented as Sup. Fig. 5B, C.

### High-throughput screening (HTS)

Cells were plated in 10 cm dishes (Corning) in full DMEM/F12 media until they reached confluence and then the media was changed to Nic media for at least 1 week. Cells were then trypsinized and plated in 30 DL at 9,000 cells/well in 384-well clear bottom white-walled plates (Greiner, Monroe, NC) in Nic media and allowed to attach overnight. Twenty-four hours after plating, media was changed (again with Nic media) and cells were treated with small molecules (Echo Liquid Handler, Beckman Coulter, Brea, CA) at a final concentration of 10 DM in DMSO. Compounds were incubated with the cells for 48 h, followed by a whole-well lytic HiBiT assay (Perkin-Elmer, Waltham, MA). A Z-factor was not calculated since no HTRA1 enhancing compounds had been identified as possible positive controls. Instead, we relied on deviation from median values across all wells within a plate. Specifically, hit compounds were defined as being > 3 standard deviations of the median plate value in the whole-well HiBiT assay. Hit compounds were then cherry-picked from a fresh source plate and verified in dose response before being repurchased for follow-up validation. HTS data are presented as [(compound signal – median signal of all compounds)/median signal of all compounds]*100. In follow up assays, we strongly recommend using compounds with minimal freeze/thaw cycles and made as fresh as possible.

### Compounds

Chicago Sky Blue 6B (CSB) was purchased from Tocris (cat. # 0846/1G), Brilliant Yellow (Bril Y) was purchased from Spectrum Chemical (cat. #BR125), Congo Red (Congo R) was purchased from Sigma Aldrich (cat. # 75768-25MG), as were Orange G (cat. # 861286-25G), Orange 2 (cat. # 195235-25G), Amaranth (cat. # A1016-50G), and Cefixime (cat. # CDS021590-100MG). Fast Green (Fast G) was purchased from Fisher (cat. # BP123-10), Bromocriptine mesylate (Bromo M) was purchased from Enzo Life Sciences (cat. # BMLD-102-0100).

### Quantitative PCR (qPCR)

Cells treated with either DMSO or CSB for 3 days were harvested by trypsinization, washed, pelleted and flash frozen before mRNA extraction (Aurum Total RNA Mini Kit, BioRad, Hercules, CA). RNA was normalized and converted to cDNA in a 5 µL reaction (qScript SuperMix, QuantaBio, Beverly, MA). cDNA was diluted 20-fold to a final volume of 100 µL with MBG water prior to qPCR analysis. TaqMan probes (Life Technologies) were used to measure *HTRA1* transcripts (Hs01016151_m1), using D*-actin* (Hs01060665_g1) as a housekeeping gene. qPCR was run on a QuantStudio 6 Real-Time qPCR system (Life Technologies) and analyzed by the accompanying software.

### Follow up HiBiT assays

The Nano-Glo HiBiT Lytic Detection System kit (Promega) was used to assess both extracellular and intracellular HiBiT levels. ARPE19 HTRA1-HiBiT or HiBiT CFH cells (cultured in either DMEM/F12 full media or Nic media, as indicated in the respective figure legends) were seeded onto 24-, 48-, or 96-well plates at a density of 200,000 cells/mL in volumes of 500, 250, or 100 DL, respectively and incubated overnight. All compounds were dissolved in DMSO as a 20 mM stock as freshly as possible. For extracellular HiBiT assays, 25 DL of media was removed from the wells and reacted with 25 DL of lytic buffer containing LgBiT (1:100) and lytic substrate (1:50), mixed for 5 min at RT and read on a GloMax Discovery (Promega). For intracellular HiBiT signal, wells were washed with PBS or HBSS followed by lysis in 50-100 DL of lytic buffer mixture (with LgBiT and substrate) diluted 1:1 with PBS or HBSS. Plates were mixed for 5 min at RT, and, if necessary (i.e., if samples were in a clear walled culture plate), samples were removed and transferred to a black walled plate prior to the luminescence reading.

### Serine hydrolase activity-based protein profiling (SH-ABPP)

For HTRA1 activity, twenty micrograms of recombinant, truncated (Gly156-Pro480) human HTRA1 protein (R&D Systems, cat. #: 2916-SE) was reconstituted in 50 mM Tris-HCl, pH 8 to make a 1.6 DM stock. Each compound (CSB, Congo R, and Bril Y) was diluted to 40 DM using the same Tris-HCl buffer.

The HTRA1 protein was mixed with the compounds at a 1:1 ratio in a final volume of 20 DL, yielding a final concentration of 800 nM protein to 20 DM compound. Samples were treated for 1 h while rotating end-over-end at RT. Denatured protein (95°C for 5 min) was used as a negative control. Samples were next reacted with 2 DM ActivX TAMRA-FP Serine Hydrolase Probe (ThermoFisher Scientific) for 1h at RT. The samples were then boiled in SDS-PAGE reducing buffer (95°C for 5 min) and run on a 4-20% Tris Gly SDS-PAGE gel. The gel was then fixed, rehydrated and imaged using the TAMRA or Cy3 channel on a Typoon Imager (GE Healthcare, Chicago, IL). Gels were then stained with Coomassie Brilliant Blue to visualize total protein and scanned electronically on an Odyssey CLx (LICOR) using the 700 nm channel. Full gel images of the recombinant HTRA1 gel are presented as Sup. Fig. 5D, E.

For whole cell-based, SH-ABPP, ARPE-19 cells expressing endogenous HTRA1 HiBiT were seeded in a 12-well format at 200,000 cells/well. Cells were then treated with the three azo dyes (CSB, Congo R, Bril Y, 20 µM, 72 h), with DMSO serving as the control (n=3 wells/treatment). Following treatment, cells were washed twice with HBSS, harvested by trypsinization, pelleted, and washed again in ice-cold HBSS. Cell pellets were lysed (25 mM Tris-HCl pH 7.4, 150 mM NaCl, 1 mM EDTA, 1% NP-40, 5% glycerol) and total protein quantification was assessed using a BCA assay. Samples were normalized to 40 Dg and reacted with 2 µM TAMRA-FP probe (1 h, RT) followed by neutralization by boiling in reducing Laemmli buffer (5 min). Samples were resolved on a 4-20% SDS-PAGE gel, fixed, and imaged on a Typhoon FLA 9500 (GE Healthcare) for FP-TAMRA signal followed by staining for total protein with Coomassie Brilliant Blue (Odyssey CLx, 700 nm channel). The area under the curve was quantified for each treatment using FIJI ImageJ, averaged and presented relative to the DMSO control.

### ARPE-19 HTRA1 stable cell line generation

A pcDNA3 construct encoding for WT or R166C^1^ human HTRA1 HiBiT with a C-terminal 6xHis tag was designed and synthesized (GenScript, Piscataway, NJ). This plasmid has a G418 selectable antibiotic marker for stable cell selection.

Eight-hundred thousand ARPE-19 cells were electroporated with 1 Dg of pcDNA plasmid using the following parameters: 1400V, 20 ms, 2 pulses. Cells were plated into antibiotic-free media in a 10 cm dish, followed by replacement with full DMEM/F12 media the next day. Stable cells that had successfully incorporated the plasmid randomly in their genome were selected using 400 Dg/mL active G418 (Sigma) for at least 2 weeks. Surviving cells were expanded and confirmed to still produce WT HTRA1 HiBiT 6xHis by routinely performing a HiBiT assay. Stable cells were seeded at a density of 20,000 cells/well in 96 well plates and allowed to attach for 1-2 days. Subsequently, they were treated with the indicated azo compounds for 72 h.

### Structural modeling and compound docking

A structural model for a full-length HTRA1 trimer was built using an alpha fold prediction of the monomer structure aligned at the trimer interface of the serine protease domain from the previously determined structure (PDBID: 7SJN). The extended trimer model was then minimized using fast relax in Rosetta. It generated 50 relaxed models that minimized the disordered regions linking the structural domains and placed them in distinct orientations. The lowest-scoring relaxed trimer model was then used to identify the binding site. Using the hugging face implementation of DiffDock^42^, the modeled trimer structure was used as a template to generate 10 complexes per molecule (Chicago Sky Blue [CSB], Congo Red [Congo R], and Brilliant Yellow [Bril Y]) using the default interface arguments. First, each molecule’s complex with the highest confidence score was selected. All three molecules were arranged in the same starting orientation based on the model, resulting in binding modes localized in three binding modes: 1-PS domain between residues 165-175, 2 – PDZ domain between residues 380-395 in an “open” conformation with no additional domain interactions, and 3 – PDZ domain between residues 380-395 with additional contacts from a loop at positions 305-310 in a “closed” domain interaction. Additionally, these resultant models were truncated to recapitulate the construct used in vitro HTRA1_156-480, and a single chain of both the extended and HTRA1_156-480 models was produced. The resulting protein-ligand complexes were then used as the starting point for further model refinement using Rosetta dock (Rosetta v. 3.13) to generate 5000 structures for each complex to produce 20,000 local structures per compound. Ligand conformations were taken from 20 conformers of each compound. Local transformations of 500 cycles did the movement in the binding site in a box of 7.0A with a move distance of 0.2 and an angle of 20, followed by high-resolution docking and rescoring. The models were then evaluated based on total complex scores to select the top 1000 models per complex. These top complexes were then analyzed based on protein: ligand interface scoring and assessed for similarity of ligand positioning based on the aspect ratio by generating a 2D histogram to visualize the explored binding space in the models. The top 5 complexes for each protein and ligand pair were then visualized using PyMOL (v.3.0.4).

## RESULTS

### CRISPR-mediated tagging results in efficient insertion of a C-terminal HiBiT hemagglutinin (HA) epitope tag sequence in the HTRA1 locus

Attempts at an N-terminal HiBiT editing strategy for HTRA1 were unsuccessful, most likely due to high GC content (79.5%) in HTRA1 exon 1 and near the desired insertion site. Therefore, we designed a genome editing strategy that would append a C-terminal HiBiT HA sequence on human HTRA1 immediately prior to its stop codon in exon 9 (Fig. 1A). Editing at the C-terminal HTRA1 locus in ARPE-19 cells was found to be very efficient (Fig. 1B, ∼88%), as genomic DNA (gDNA) from unenriched heterogenous pools of cells revealed two bands, an unedited lower molecular weight band, and a higher molecular weight band corresponding to insertion of 72 bp (VS HiBiT GG HA). This high level of editing efficiency enabled us to use a heterogenous pool of HTRA1 HiBiT cells instead of isolating clonal lines, which can have wildly different growth and behavioral properties. Moreover, the top, edited gDNA band was excised and sent for Sanger sequencing to confirm proper insertion of the HiBiT HA tag (Sup. Fig. 1), which exactly matched our desired design (Fig. 1A).

**Figure 1.**
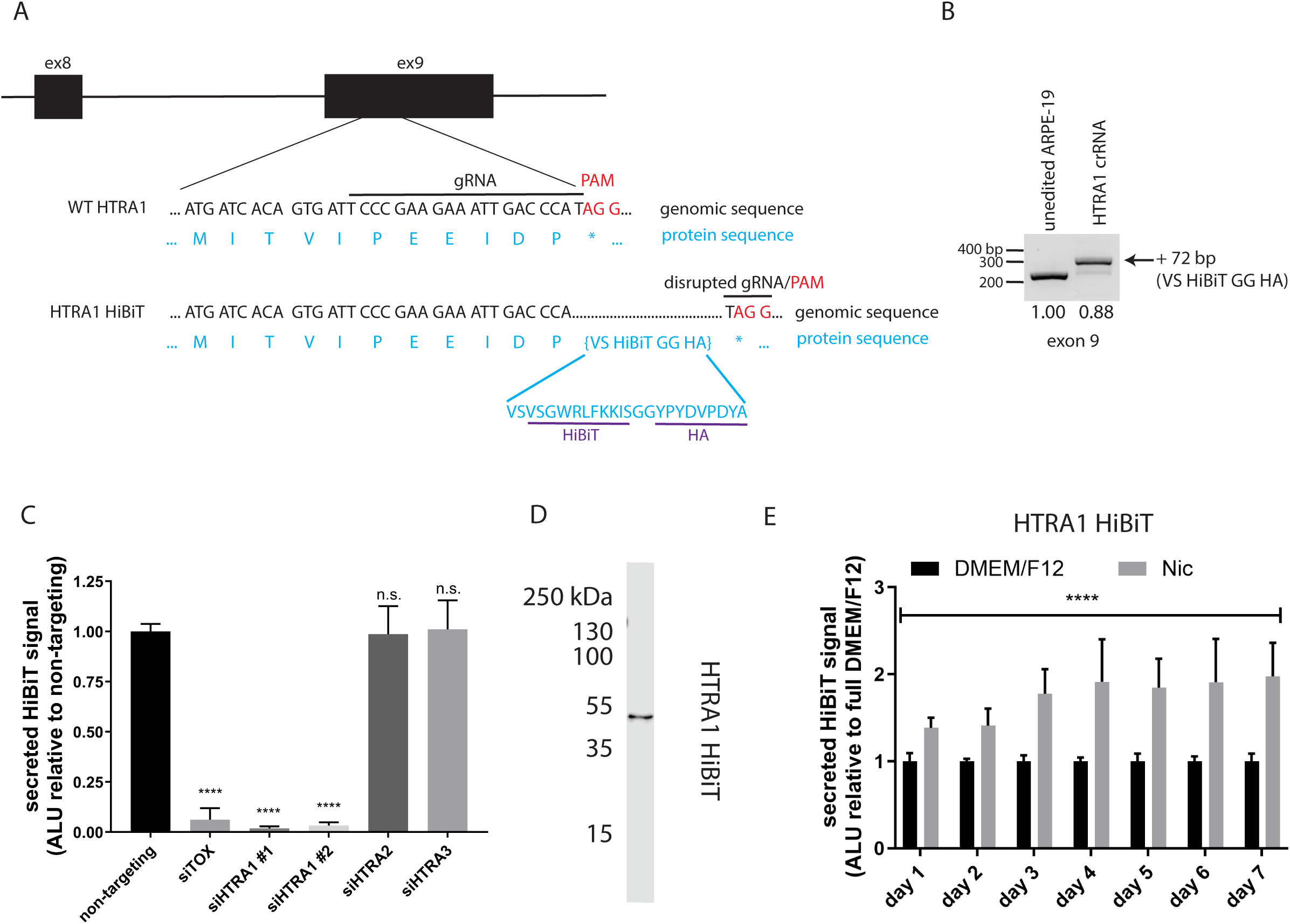
Design and validation of HTRA1 HiBiT editing strategy in ARPE-19 cells. (A) HTRA1 genomic DNA sequence and design of a C-terminal HiBiT HA insertion. (B) HTRA1 exon 9 gDNA amplification of unedited and edited ARPE-19 cells. Insertion of the VS HiBiT GG HA sequence increases the predicted molecular weight by 72 bp. Estimated editing efficiency (top band/bottom band) is ∼88%. (C) Validation of HiBiT insertion at the HTRA1 locus by siRNA. ARPE-19 HTRA1 HiBiT edited cells were reverse transfected with control siRNAs (non-targeting, siTOX), or siRNAs against HTRA1, HTRA2, or HTRA3. n = 4 independent experiments for HTRA1 siRNAs, n = 2 independent experiments for HTRA2 and HTRA3 siRNAs. n.s. = not significant, **** p < 0.0001, one-way ANOVA with multiple comparisons vs. non-targeting. (D) HTRA1 HiBiT protein migrates at the expected molecular weight of ∼55 kDa. ARPE-19 HTRA1 HiBiT cells were incubated in serum free media followed by concentration and analysis by HiBiT blotting. n = 3 independent experiments. (E) Nic differentiation media increases HTRA1 secretion. ARPE-19 HTRA1 HiBiT cells were plated at confluence followed by incubation in either full DMEM/F12 or Nic media for up to 7 days. A HiBiT assay was performed on media aliquots for 1 week. n = 3 independent experiments, **** p < 0.0001, unpaired t-test with multiple comparisons vs. respective DMEM/F12 control.

### HiBiT tagging specifically modifies HTRA1

Next, we used short interfering RNA (siRNA) to confirm that any observed HiBiT signal produced from the ARPE-19 HTRA1 HiBiT cells originated only from the editing of HTRA1, and not from homologous proteins, such as HTRA2 or HTRA3. Two independent HTRA1 siRNAs, but not HTRA2 or HTRA3 siRNAs, significantly reduced extracellular HTRA1 HiBiT signal in edited cells (97-98% reduction on average compared to non-targeting siRNA, *** p < 0.001, ANOVA, Fig. 1C). siTOX, a pool of proprietary siRNA that kills cells upon successful transfection, was used as a positive control to estimate transfection efficiency during each reverse transfection. After validating the specificity of the HiBiT tagging, we also confirmed that the resultant endogenous HTRA1 HiBiT protein produced matched apparent molecular weight and isoform expectations. Indeed, secreted HTRA1 HiBiT appeared as a single species at ∼50 kDa under reducing and denaturing conditions, in accordance with expectations (https://www.uniprot.org/). Finally, since we edited the endogenous HTRA1 locus, we also tested whether the standard culture conditions used for ARPE-19 cells (i.e., DMEM/F12 in 10% FBS) were ideal for producing the HTRA1 protein. While culturing in DMEM/F12 full media resulted in sufficient levels of HTRA1 HiBiT for easy and sensitive detection, we found that differentiation conditions using media low in serum (1% FBS) and containing 10 mM nicotinamide^40^ significantly increased HTRA1 secreted protein levels by up to 2-fold (Fig. 1E, **** p < 0.001, t-test with multiple comparisons vs. DMEM/F12 full media). Thus, we used these RPE differentiation conditions for performing high-throughput screening (HTS) to identify HTRA1 enhancers (see below).

### HTS of HTRA1 HiBiT cells identifies Chicago sky blue 6B (CSB) as an enhancing molecule

While the HiBiT tag has been used to perform a series of screens designed to sensitively identify small molecule degraders of endogenous POIs^43, 44^, we envisioned that the HiBiT tag would also enable identification of small molecules which act as chemical chaperones, transcriptional enhancers, or protein stabilizers. In theory, such molecules could serve as therapeutics to increase HTRA1 levels and activity in individuals with HTRA1-based AMD risk alleles (i.e., rs362127733^15^) or may serve as chemical probes/lead compounds for future development. To identify molecules with such properties, we screened ∼1920 annotated compounds from chemical libraries (originating from the National Institutes of Health Clinical and Prestwick Collections). A Z factor^45^, a typical metric for HTS, was not possible since no known compounds have been identified that enhance HTRA1 levels. Therefore, we used three standard deviations above the plate median to identify HTRA1 enhancing compounds (Fig. 2A).

**Figure 2.**
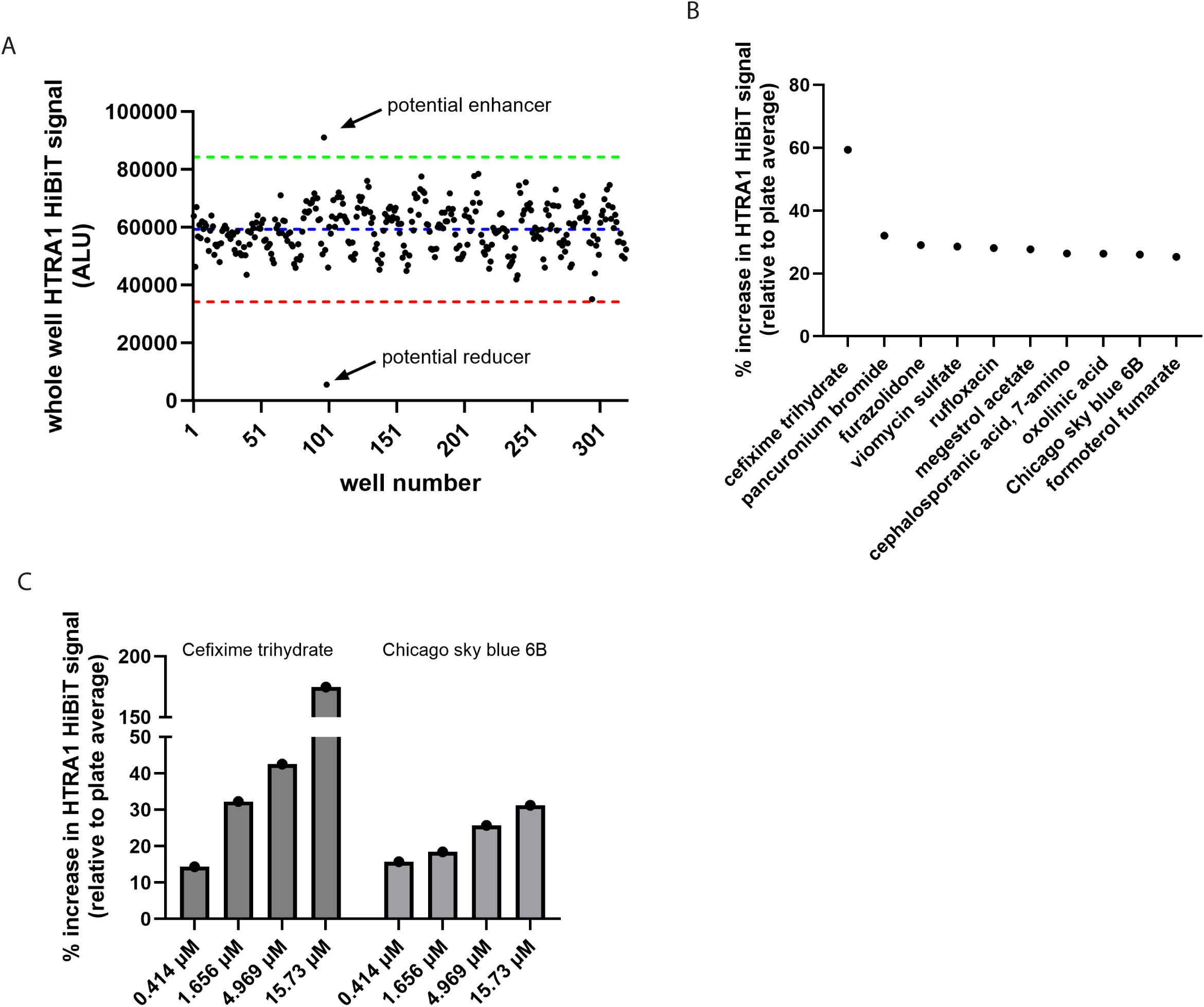
Summary of HTS primary screen and dose-response. (A) An example HTS assay plate from the primary screen of the NIH and Prestwick libraries. Three standard deviations of the mean is shown by the dotted green line whereas three standard deviations below the mean is shown by the dashed red line. Compounds outside of these lines were considered hit compounds. (B) Top ten compounds from a list of 17 small molecules identified in the primary screen. (C) Reproduced dose-responsiveness of two hit compounds, cefixime trihydrate and Chicago Sky Blue 6B.

ARPE-19 HTRA1 HiBiT cells were differentiated at confluence in Nic media for at least 1 week prior to replating and treatment with 10 DM small molecule for 48 h. This treatment was followed by a whole-well (combined extracellular and intracellular HTRA1) HiBiT assay using HiBiT lytic reagent. Seventeen compounds were identified as enhancers, resulting in a hit rate of 0.89%. The initial top ten hit compounds identified in the primary screen included (in descending order): cefixime trihydrate, pancuronium bromide, furazolidone, viomycin sulfate, rufloxacin, megestrol acetate, 7-amino-cephalosporanic acid, oxolinic acid, CSB, and formoterol fumarate (Fig. 2B). Of the originally identified hits, after cherry picking, only cefixime trihydrate and CSB reproduced in dose-response studies (Fig. 2C). Unfortunately, while cefixime appeared to be the most promising hit, this compound list was further whittled to only CSB (Fig. 3A) upon repurchasing the compound and deconvoluting the previous whole-well assay into separate extracellular and intracellular assays (Fig. 3B). CSB demonstrated a dose-responsive effect, significantly increasing extracellular and intracellular HTRA1 HiBiT at 20 DM (Fig. 3B, 1.7 ± 0.2 fold, and 2.0 ± 0.3 fold, respectively, ** p < 0.001, *** p < 0.0001, ANOVA). These HiBiT assay results were also verified at the protein level using HiBiT blotting (Fig. 3C).

**Figure 3.**
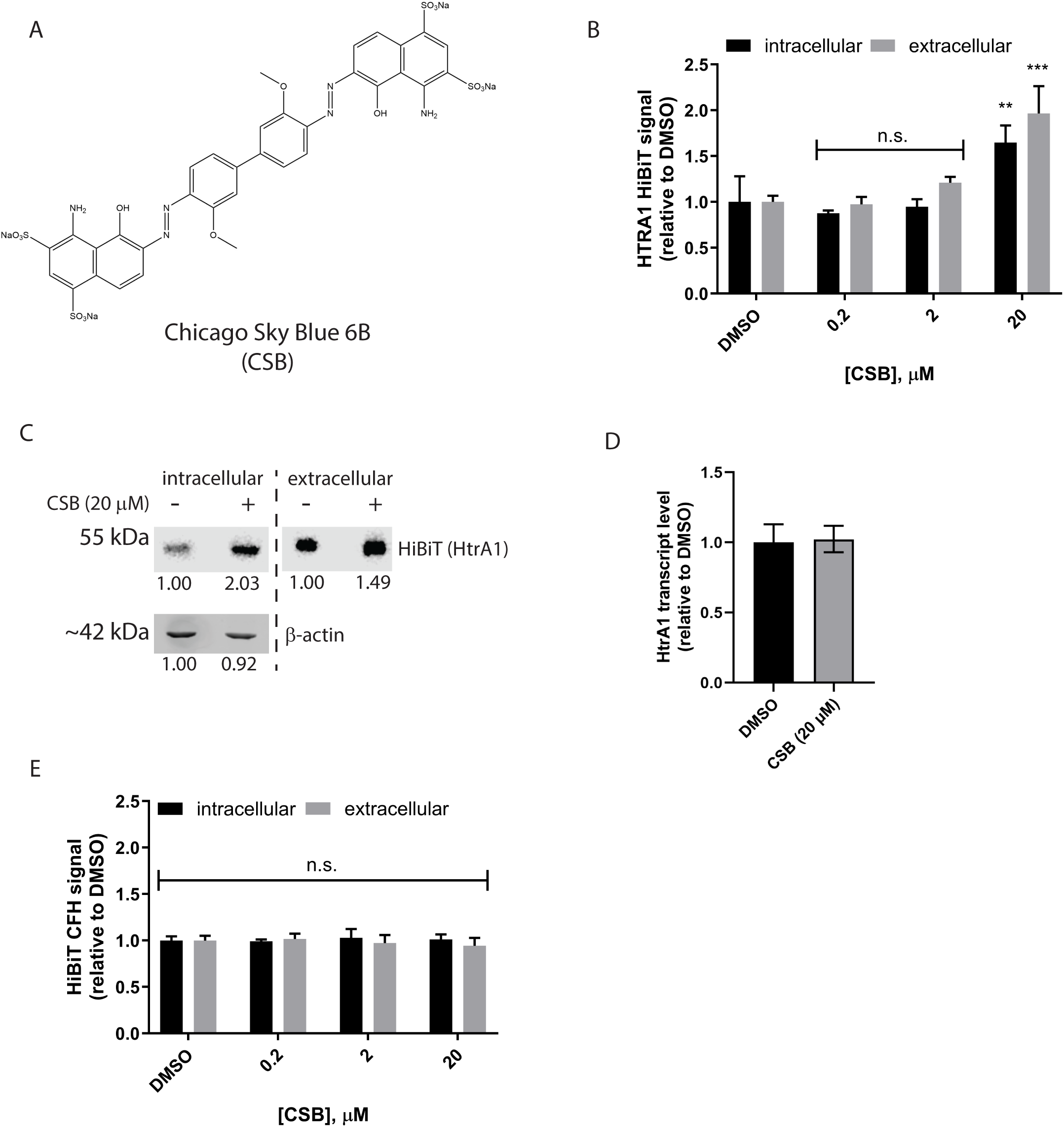
Confirmatory studies on the effect of Chicago Sky Blue 6B (CSB) on HTRA1 HiBiT produced from ARPE-19 cells. (A) Azo-based chemical structure of CSB. (B) Validation of the effects of CSB and deconvolution of the previous whole well results into extracellular and intracellular components. ARPE-19 HTRA1 HiBiT cells (grown in Nic media) were treated with the indicated concentration of CSB for 72 h. n = 3-4 independent experiments, n.s. = not significant, ** p < 0.01, *** p < 0.001, one-way ANOVA with multiple comparisons vs. DMSO. (C) CSB increases HTRA1 HiBiT protein levels intracellularly and extracellularly as determined by a HiBiT assay. Cells were treated with 20 DM CSB for 72 h in serum free media followed by concentration of the conditioned media, lysis of the cells and HiBiT blotting. Representative data of three independent experiments. (D) CSB does not alter HTRA1 expression levels. ARPE-19 cells were treated with 20 DM CSB for 72 h in Nic media followed by mRNA extraction and qPCR. Representative data of three independent experiments. (E) CSB has no effect on HiBiT CFH secretion or intracellular levels. A control ARPE-19 cell line expressing HiBiT CFH (Sup. Fig. 2) was treated with CSB at the indicated concentration for 72 h, followed by HiBiT assay. N = 3 independent experiments n.s. = not significant, one-way ANOVA with multiple comparisons vs. DMSO.

While we sought to identify compounds that increased HTRA1, possibly by increased transcription, at this point, it was unclear how CSB was increasing HTRA1 HiBiT levels. Surprisingly, quantitative PCR (qPCR) demonstrated that CSB did not increase HTRA1 transcript levels (Fig. 3D). These results suggest that CSB is acting at the post-transcriptional level, potentially through HTRA1 protein stabilization.

To apply a degree of specificity to our findings, we generated an additional CRISPR-engineered cell line, this time expressing HiBiT complement factor H (CFH). The design and validation of this line is presented in Sup. Fig. 2 and Sup. Fig. 3. This CFH control cell line allows us to decipher whether the observed effects of CSB were potentially due to the HiBiT tag itself, or if CSB increases the levels of multiple secreted proteins in a more general sense. However, CSB had no effect on CFH extracellular or intracellular levels (Fig. 3E) under the same conditions used for HTRA1 HiBiT cells in Fig. 3B.

### Congo Red, an azo dye with structural similarities to CSB also regulates HTRA1 abundance

CSB belongs to a family of compounds called azo dyes, which are characterized by two R-N=N-R’ groups. These compounds are typically brightly colored and used in food, textiles, or as stains^46^. To determine if the effects we observed on HTRA1 were due to the unique structure of CSB, or if they were, perhaps, more broadly representative of azo dyes in general, we tested the effects of Congo Red (Congo R, Fig. 4A), typically used as an amyloid stain^47^, and Brilliant Yellow (Bril Y, Fig. 4B), a textile dye, on HTRA1 levels. At a very basic level, while Congo R mimics the aromatic base structure of CSB (cf. Fig. 3A to Fig. 4A), Bril Y is more distinct from CSB in its chemical structure and spacing. Intriguingly, Congo R (20 DM, 72 h) had profound elevating effects on intracellular endogenous HTRA1 HiBiT (3.6 ± 0.3, *** p < 0.001, ANOVA), but not on secreted levels (Fig. 4C), whereas Bril Y (also 20 DM, 72 h) had no effect on either metric (Fig. 4C). HTRA1 extracellular and intracellular levels were also unaffected by a series of monoazo dyes, including Orange G, Orange 2, Amaranth, as well as a non-azo dye, Fast Green (Sup. Fig. 4A-D, F). Finally, we also tested bromocriptine mesylate as a potential functional mimetic (VGLUT1 inhibitor^48^) of CSB, which also had no effect on HTRA1 secretion or intracellular levels (Sup. Fig. 4E, F). These results suggest that perhaps the symmetry and structure of the azo dyes, CSB and Congo R, are necessary components for HTRA1 enhancement.

**Figure 4.**
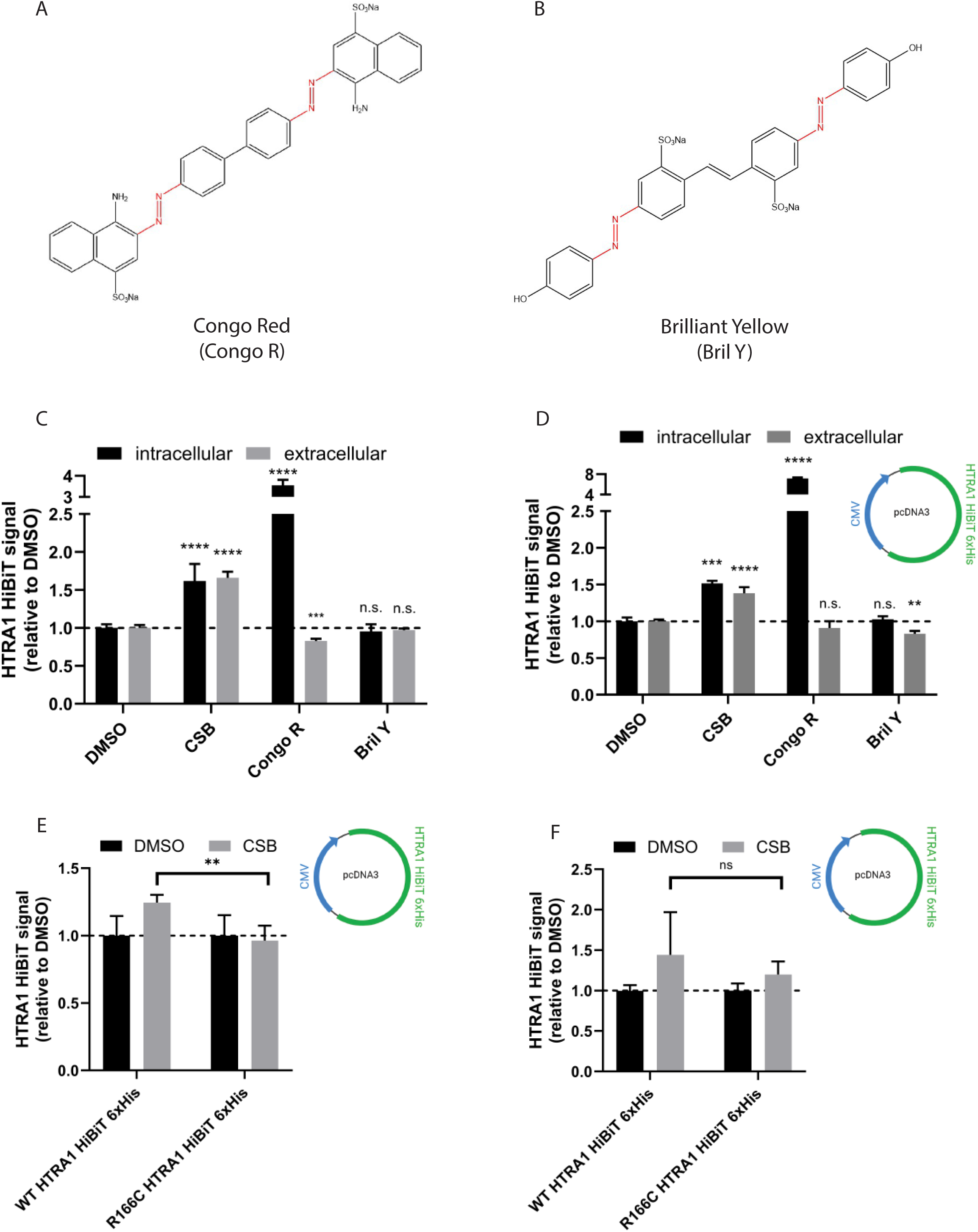
Additional azo dyes affect HTRA1 levels. (A, B) Chemical structures of azo dyes Congo Red (Congo R) and Brilliant Yellow (Bril Y). (C) Congo R, but not Bril Y increases HTRA1 intracellular levels. ARPE-19 HTRA1 HiBiT cells grown in DMEM/F12 full media were treated with the indicated compound (20 DM) for 72 h followed by a HiBiT assay. Representative data of 3 independent experiments. n.s. = not significant, *** p < 0.001, **** p < 0.0001, one-way ANOVA with multiple comparisons vs. DMSO control. Note: Congo R data were compared to a different DMSO control than presented. (D) Azo dyes similarly alter CMV-driven HTRA1 HiBiT levels. ARPE-19 cells expressing an overexpression plasmid encoding for HTRA1 HiBiT 6xHis were treated as described in (C) followed by a HiBiT assay. Representative data of 3 independent experiments. n.s. = not significant, ** p < 0.01, *** p < 0.001, **** p < 0.0001, one-way ANOVA with multiple comparisons vs. DMSO control. (E, F) HTRA1 variants with mutations near the dimer interface are affected by CSB differently than WT HTRA1. (E) Stable ARPE-19 cells expressing WT or R166C HTRA1 HiBiT 6xHis were treated with CSB for 72 h followed by a HiBiT assay of (E) extracellular or (F) intracellular HTRA1. Data pooled from three independent experiments. n.s. = not significant, ** p < 0.01, two sided t-test.

To further confirm that these enhancing effects were indeed post-translational, we engineered a new ARPE-19 cell line that, instead of incorporation of HiBiT into the endogenous HTRA1 locus, used a plasmid-based, CMV-driven wild-type HTRA1 HiBiT 6xHis construct. Thus, the expression of this version of HTRA1 would be controlled by completely different transcriptional regulatory elements than our original ARPE-19 (endogenous) HTRA1 HiBiT cell line. After treatment of this HTRA1 overexpression cell line with CSB, Congo R, or Bril Y (20 DM, 72 h), we found a striking degree of parallel findings to the treatment of endogenously-regulated HTRA1 HiBiT ARPE-19 cells. CSB significantly increased extracellular (Fig. 4D, 1.4 ± 0.1 fold, *** p < 0.001, ANOVA) and intracellular HTRA1 HiBiT 6xHis (Fig. 4D, 1.5 ± 0.03, *** p < 0.001, ANOVA). Congo R elevated only HTRA1 HiBiT 6xHis intracellular levels, but to extreme levels (Fig. 4D, 7.1 ± 0.2 fold, *** p <0.001, ANOVA), whereas Bril Y did not raise HTRA1 HiBiT 6xHis levels (Fig. 4D). These results reaffirm the transcriptional independence of our azo-mediated HTRA1 observations.

There is a consensus in the scientific community that HTRA1 exists in a dynamic equilibrium between monomers and trimer^2^, with the trimers being the more stable of the two species^3, 4^. Thus, compounds that promote trimerization may also increase HTRA1 levels through reducing HTRA1 turnover. Thus, we speculated that CSB was capable of increasing HTRA1 levels by stabilizing the trimer, thereby increasing the HiBiT signal. To partially test this idea, we generated an R166C variant of HTRA1 which has a disrupted trimer interface^1^. Interestingly, CSB did not increase R166C HTRA1 extracellular levels compared to WT HTRA1 (Fig. 4E), and minimized corresponding CSB-mediated increases intracellularly, albeit not to a significant extent (Fig. 4F, p = 0.11). These results hint that CSB may be working via stabilizing the HTRA1 trimer interface.

### Congo R, but not CSB inhibits HTRA1 enzymatic activity

As a secreted serine hydrolase, HTRA1 abundance, as determined by a HiBiT assay or blot does not necessarily indicate that the protein is active. Activity of an enzyme will depend on its degree of foldedness, processing (if applicable), and the presence (or absence) of enzyme inhibitors or cofactors^49^. Therefore, we also tested whether azo compounds had any impact on recombinant, active human HTRA1 using a chemical biology approach called serine hydrolase activity-based protein profiling (SH-ABPP)^50, 51^ which results in covalent fluorescent modification (TAMRA) of enzymes if they are active and can react with the generalized fluorophosphonate (FP) probe. Treatment of recombinant HTRA1 with 20 DM CSB, Congo R, or Bril Y for 1 h demonstrated that CSB and Bril Y had no effect on HTRA1 activity (Fig. 5A, B) relative to DMSO treatment. However, surprisingly, Congo R significantly decreased recombinant HTRA1 activity by over half (Fig. 5A, B, 0.45 ± 0.07 fold, *** p < 0.001, ANOVA). These results suggest that some azo compounds can directly bind to HTRA1 and affect its activity by a yet-to-be determined mechanism.

**Figure 5.**
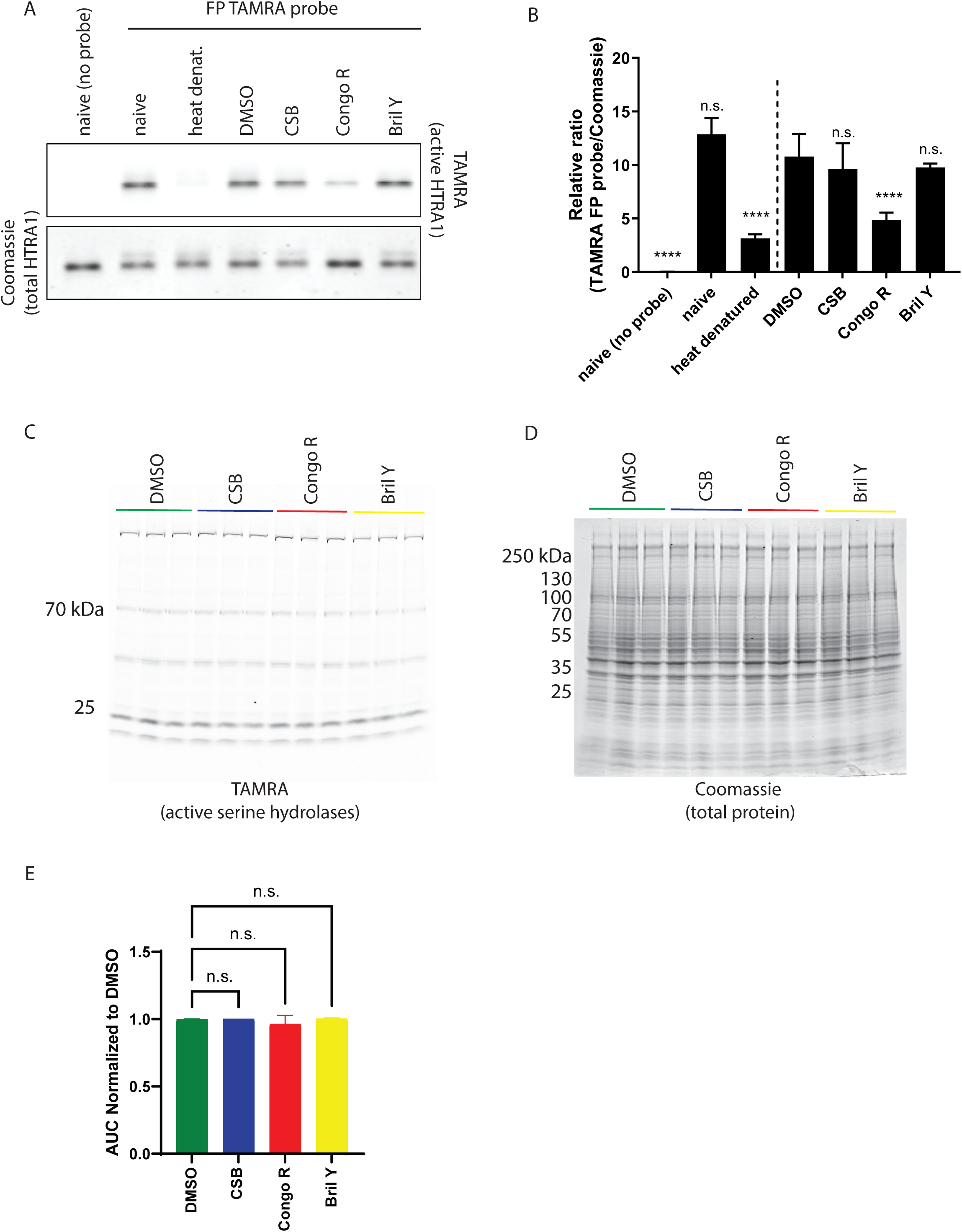
Serine hydrolase activity-based protein profiling (SH-ABPP) post azo compound treatment. (A) Congo R inhibits human recombinant HTRA1 protein activity. HTRA1 protein was incubated with 20 mM of compound for 1 h followed by SH-ABPP using a FP-TAMRA probe. (B) FP-TAMRA signal was normalized to total protein (Coomassie). Representative data shown of at least two independent experiments performed in technical duplicate/triplicate. n.s. = not significant, **** p < 0.0001, one-way ANOVA with multiple comparison test vs. DMSO control. (C, D) Whole cell SH-APBB of ARPE-19 (endogenous) HTRA1 HiBiT cells treated with azo dyes (20 DM, 72 h) and the corresponding total protein stain (Coomassie). Representative data of three independent experiments. (E) Quantification of total TAMRA signal across all experiments and normalized to DMSO controls. n.s. = not significant, one-way ANOVA.

### Azo compounds do not broadly alter the activity of other cellular serine hydrolases

We next asked whether the same azo compounds (CSB, Congo R, Bril Y) had any inhibitory effect on other serine hydrolases with similar active sites^52^ in whole cell lysates. ARPE-19 cells expressing endogenous HTRA1 HiBiT were treated for 72 h with 20 DM compound followed by cell harvesting, lysis, and reaction with a FP TAMRA similarly to that described above for recombinant HTRA1. Intriguingly, we found that none of the intracellular serine hydrolases were affected by any of the azo dyes (Fig. 5C, D). Additionally, total protein content as determined by Coomassie Blue, demonstrated that these dyes also do not globally change cellular protein abundance (Fig. 5E).

### Azo compounds potentially interact with HTRA1 at its trimer interface

HTRA1 trimerizes at the trypsin-like serine protease (SP) domain and is flanked by the N-terminal IGFBP and Kazal-like domains and the C-terminal PDZ domain. To identify potential binding modes of the compounds to HTRA1, we performed an unbiased search for docking sites of the three azo compounds (CSB, Congo R, and Bril Y) on a full-length model of HTRA1 using DiffDock^53^. Then, we assessed the differences in binding modes across the three compounds using docking in Rosetta (Fig. 6A). The first modeling stage in DiffDock identified different surfaces between the azo compounds, with higher confidence docking for Bril Y and Congo R than CSB (Sup. Fig. 6A-D). However, all three compounds had putative binding regions at the non-catalytic face of the trimerized serine protease domain (Fig. 6B-D). To assess the differences in binding at the timer interface among the three compounds, we evaluated the relative movement of the compounds from the initial DiffDock binding pose using docking. Surprisingly, we found that CSB appears to have the lowest aspect ratio while having a higher interface score, indicating a weaker, but more defined binding ability that was closer to the initial binding mode (Fig. 6B). Congo R, on the other hand, demonstrated a higher aspect ratio and lower interface score, indicating stronger binding, but with increased movement of the molecule with respect to initial binding site (Fig. 6C). Bril Y demonstrated both a high aspect ratio and high interface score, suggesting an undefined interaction potential (Fig. 6D). Basic (H167 and K168) and aromatic (Y169) residues on HTRA1 mediate key contacts to sulfonate groups and the ring structures of the ligands (Fig. 6E-G). Interestingly, comparing cumulative contacts for the ensembles shows that the defined Congo R contacts may be a result of asymmetry of binding to the trimer shown by interactions dominated by one chain while contacts to CSB and Bril Y appear to be distributed. These computational findings may ultimately provide clues as to how azo dyes may interact with and modify HTRA1 stability, trimerization, and function.

**Figure 6.**
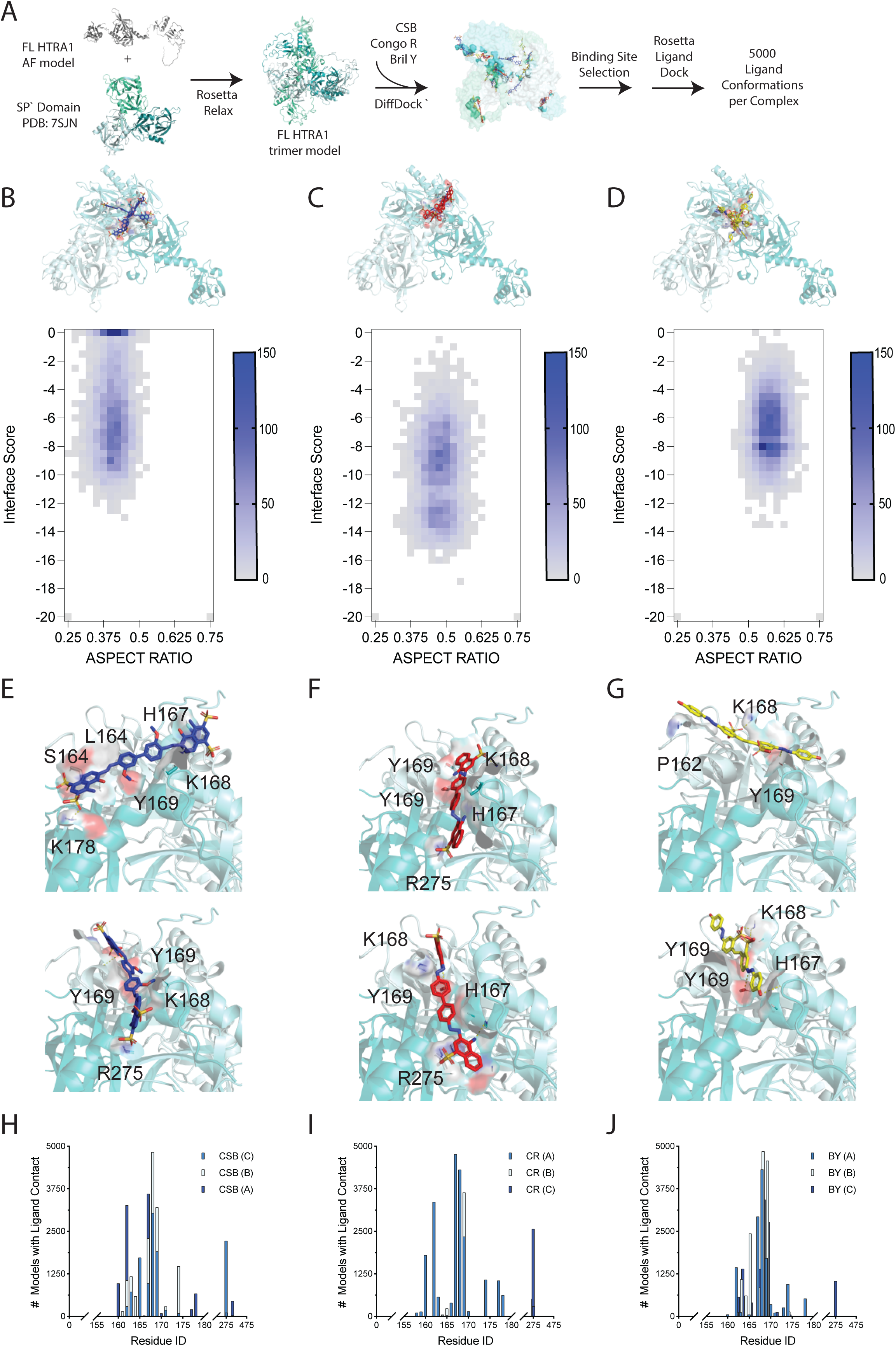
Compound binding to HTRA1 simulation. (A) Workflow for modeling interactions between azo compounds and HTRA1 models. (B-D) Binding site models of the top 5 scoring ligand positions, surface representation (top). 2D histogram of the sampled ligand positions (aspect ratio) and interface scores for CSB (B) Congo R (C) and Bril Y (D) colored from gray to blue based on the sampling frequency (bottom). A lower aspect ratio indicates less tolerated movements of the molecule within the model, whereas a higher interface score indicates Binding site models highlighting interactions between ligand the trimeric HTRA1 and CSB (E), Congo R (F) and Bril Y (G). HTRA is shown in cartoon representation. Interacting residues are shown in surface and stick representation. CSB, Congo R and Bril Y are shown in sticks and colored blue, red and yellow, respectively. Residues interacting in each model are labelled. (H-J) Summary of cumulative ligand residue contacts across modeled ensembles between HTRA1 and CSB (H), Congo R (I) and Bril Y(J). Residue contacts to ligand are shown for each HTRA1 chain in the trimer defined as A, B and C.

## DISCUSSION

CSB and derivatives thereof have been used as vital dyes since the 1930’s to assist with vascular imaging^54^ and intrathoracic surgeries^55, 56^. CSB itself has since been demonstrated to have bioactive functions in culture and in vivo including promoting cardiac repair, neuroprotection, and prevention of amyloid formation^48, 57–61^, highlighting its diverse capabilities outside of its use as a dye/imaging agent. Using unbiased screening, we identified CSB as a small molecule capable of significantly elevating HTRA1 levels in culture while not affecting its activity, adding to this molecule’s known roles. It is intriguing to speculate that perhaps some of the previously identified capabilities of CSB on amyloid formation or neuroprotection, for example, could be due in part to CSB-mediated increases in HTRA1^62, 63^.

Our findings have potential implications for diseases affected by alterations in HTRA1 activity, either due to transcriptional alterations or mutations. For example, high levels of CSB would be predicted to increase HTRA1 activity by ∼2-fold through promoting its stability and/or decreasing its turnover. Theoretically, such a molecule may be effective in partially counteracting age-dependent decreases (∼2.8 fold) in RPE-choroid HTRA1 protein observed in AMD risk donors with the rs10490924 SNP^15^. CSB could also serve as a lead molecule for other non-AMD loss-of-function diseases caused by HTRA1 mutations such as cerebral autosomal-recessive arteriopathy with subcortical infarcts and leukoencephalopathy (CARASIL)^64^. While there are many instances that null HTRA1 mutations cause CARASIL, select loss-of-function mutations identified in symptomatic carriers or CARASIL patients such as A173P, P258Q, F286V, R166C, A173T, A321T, and L364P still retain partial activity^65^. Thus, molecules like CSB or CSB derivatives may be useful for recovering lost HTRA1 activity in these instances, potentially altering CARASIL severity or disease trajectory.

Our computational modeling work suggests that CSB and Congo R have distinct interactions at the HTRA1 trimer interface. After studying the small molecule docking findings, it is possible that the charged sulfate groups in CSB may be key in mediating its modulatory activity, elevating steady state levels of the protein while not affecting its enzymatic activity. Conversely, due to its overall higher nonpolar aromatic properties, Congo R may bind with “higher” affinity to HTRA1, but in less specific binding modes which appear to influence biological activity. These data present new ideas for how this group of molecules can be optimized to improve specificity by placement of charged groups that may help orient molecules in defined binding modes.

Of course, there are also challenges associated with the use of azo dyes, including potential environmental concerns such as bioaccumulation, prolonged environmental stability, and inefficient biodegradation mechanisms^66^. Moreover, significant improvement to the potency of CSB must be made for it to be effective in vivo, potentially within the eye. Nonetheless, our study provides a blueprint for sensitively monitoring both HTRA1 and CFH through HiBiT tagging, and our findings identify a previously undiscovered tool compound (CSB) upon which researchers can build and manipulate in the hopes of identifying effective and safe HTRA1 enhancing molecules for disease treatment.

## ACKNOWLEDGEMENTS

JDH is the Larson Endowed Chair for Macular Degeneration Research (UMN). The work described herein was supported by the Helen Lindsay Family Foundation and R21 EY032693 (JDH). JDH received support previously from an endowment from the Roger and Dorothy Hirl Research Fund, a Macular Degeneration Research Grant from the Fichtenbaum Charitable Trust, and P30 EY030413 (to the UT Southwestern Department of Ophthalmology). JDH is also a member of the Promega Advanced Academic Access Program, which provided reagent support for high-throughput screening. SB was supported by an NIH F31 grant from NINDS (F31 NS12751301). AA was supported by an NIH T35 Training Grant (T35 EY026510). LAJ is supported by an Effie Marie Cain Scholarship in Medical Research and by an NIH grant (R01 AG076459).

## Notes

### Competing Interest Statement

The authors have declared no competing interest.

## REFERENCES

(1) Springelkamp, H.; Mishra, A.; Hysi, P. G.; Gharahkhani, P.; Hohn, R.; Khor, C. C.; Cooke Bailey, J. N.; Luo, X.; Ramdas, W. D.; Vithana, E.; et al. Meta-analysis of Genome-Wide Association Studies Identifies Novel Loci Associated With Optic Disc Morphology. Genetic epidemiology 2015, 39 (3), 207–216. DOI: 10.1002/gepi.21886.

(2) Cheng, C. Y.; Schache, M.; Ikram, M. K.; Young, T. L.; Guggenheim, J. A.; Vitart, V.; MacGregor, S.; Verhoeven, V. J.; Barathi, V. A.; Liao, J.; et al. Nine loci for ocular axial length identified through genome-wide association studies, including shared loci with refractive error. American journal of human genetics 2013, 93 (2), 264–277. DOI: 10.1016/j.ajhg.2013.06.016.

(3) Gharahkhani, P.; Jorgenson, E.; Hysi, P.; Khawaja, A. P.; Pendergrass, S.; Han, X.; Ong, J. S.; Hewitt, A. W.; Segre, A. V.; Rouhana, J. M.; et al. Genome-wide meta-analysis identifies 127 open-angle glaucoma loci with consistent effect across ancestries. Nature communications 2021, 12 (1), 1258. DOI: 10.1038/s41467-020-20851-4.

(4) Fritsche, L. G.; Igl, W.; Bailey, J. N.; Grassmann, F.; Sengupta, S.; Bragg-Gresham, J. L.; Burdon, K. P.; Hebbring, S. J.; Wen, C.; Gorski, M.; et al. A large genome-wide association study of age-related macular degeneration highlights contributions of rare and common variants. Nature genetics 2016, 48 (2), 134–143. DOI: 10.1038/ng.3448.

(5) Winkler, T. W.; Grassmann, F.; Brandl, C.; Kiel, C.; Gunther, F.; Strunz, T.; Weidner, L.; Zimmermann, M. E.; Korb, C. A.; Poplawski, A.; et al. Genome-wide association meta-analysis for early age-related macular degeneration highlights novel loci and insights for advanced disease. BMC Med Genomics 2020, 13 (1), 120. DOI: 10.1186/s12920-020-00760-7.

(6) Wong, W. L.; Su, X.; Li, X.; Cheung, C. M.; Klein, R.; Cheng, C. Y.; Wong, T. Y. Global prevalence of age-related macular degeneration and disease burden projection for 2020 and 2040: a systematic review and meta-analysis. The Lancet. Global health 2014, 2 (2), e106–116. DOI: 10.1016/S2214-109X(13)70145-1.

(7) Fleckenstein, M.; Keenan, T. D. L.; Guymer, R. H.; Chakravarthy, U.; Schmitz-Valckenberg, S.; Klaver, C. C.; Wong, W. T.; Chew, E. Y. Age-related macular degeneration. Nat Rev Dis Primers 2021, 7 (1), 31. DOI: 10.1038/s41572-021-00265-2.

(8) Fritsche, L. G.; Fariss, R. N.; Stambolian, D.; Abecasis, G. R.; Curcio, C. A.; Swaroop, A. Age-related macular degeneration: genetics and biology coming together. Annu Rev Genomics Hum Genet 2014, 15, 151–171. DOI: 10.1146/annurev-genom-090413-025610 From NLM Medline.

(9) Pan, Y.; Fu, Y.; Baird, P. N.; Guymer, R. H.; Das, T.; Iwata, T. Exploring the contribution of ARMS2 and HTRA1 genetic risk factors in age-related macular degeneration. Progress in retinal and eye research 2023, 97, 101159. DOI: 10.1016/j.preteyeres.2022.101159 From NLM Medline.

(10) Iejima, D.; Itabashi, T.; Kawamura, Y.; Noda, T.; Yuasa, S.; Fukuda, K.; Oka, C.; Iwata, T. HTRA1 (high temperature requirement A serine peptidase 1) gene is transcriptionally regulated by insertion/deletion nucleotides located at the 3’ end of the ARMS2 (age-related maculopathy susceptibility 2) gene in patients with age-related macular degeneration. The Journal of biological chemistry 2015, 290 (5), 2784–2797. DOI: 10.1074/jbc.M114.593384.

(11) Iejima, D.; Nakayama, M.; Iwata, T. HTRA1 Overexpression Induces the Exudative Form of Age-related Macular Degeneration. J Stem Cells 2015, 10 (3), 193–203.

(12) Nakayama, M.; Iejima, D.; Akahori, M.; Kamei, J.; Goto, A.; Iwata, T. Overexpression of HtrA1 and exposure to mainstream cigarette smoke leads to choroidal neovascularization and subretinal deposits in aged mice. Investigative ophthalmology & visual science 2014, 55 (10), 6514–6523. DOI: 10.1167/iovs.14-14453 From NLM Medline.

(13) Jones, A.; Kumar, S.; Zhang, N.; Tong, Z.; Yang, J. H.; Watt, C.; Anderson, J.; Amrita Fillerup, H.; McCloskey, M.; et al. Increased expression of multifunctional serine protease, HTRA1, in retinal pigment epithelium induces polypoidal choroidal vasculopathy in mice. Proceedings of the National Academy of Sciences of the United States of America 2011, 108 (35), 14578–14583. DOI: 10.1073/pnas.1102853108.

(14) Owen, L. A.; Shirer, K.; Collazo, S. A.; Szczotka, K.; Baker, S.; Wood, B.; Carroll, L.; Haaland, B.; Iwata, T.; Katikaneni, L. D.; DeAngelis, M. M. The Serine Protease HTRA-1 Is a Biomarker for ROP and Mediates Retinal Neovascularization. Front Mol Neurosci 2020, 13, 605918. DOI: 10.3389/fnmol.2020.605918 From NLM PubMed-not-MEDLINE.

(15) Williams, B. L.; Seager, N. A.; Gardiner, J. D.; Pappas, C. M.; Cronin, M. C.; Amat di San Filippo, C.; Anstadt, R. A.; Liu, J.; Toso, M. A.; Nichols, L.; et al. Chromosome 10q26-driven age-related macular degeneration is associated with reduced levels of HTRA1 in human retinal pigment epithelium. Proceedings of the National Academy of Sciences of the United States of America 2021, 118 (30). DOI: 10.1073/pnas.2103617118.

(16) Lin, M. K.; Yang, J.; Hsu, C. W.; Gore, A.; Bassuk, A. G.; Brown, L. M.; Colligan, R.; Sengillo, J. D.; Mahajan, V. B.; Tsang, S. H. HTRA1, an age-related macular degeneration protease, processes extracellular matrix proteins EFEMP1 and TSP1. Aging Cell 2018, 17 (4), e12710. DOI: 10.1111/acel.12710.

(17) Hulleman, J. D. Malattia Leventinese/Doyne Honeycomb Retinal Dystrophy: Similarities to Age-Related Macular Degeneration and Potential Therapies. Adv Exp Med Biol 2016, 854, 153–158. DOI: 10.1007/978-3-319-17121-0_21.

(18) Stone, E. M.; Braun, T. A.; Russell, S. R.; Kuehn, M. H.; Lotery, A. J.; Moore, P. A.; Eastman, C. G.; Casavant, T. L.; Sheffield, V. C. Missense variations in the fibulin 5 gene and age-related macular degeneration. The New England journal of medicine 2004, 351 (4), 346–353. DOI: 10.1056/NEJMoa040833 351/4/346 [pii].

(19) Mullins, R. F.; Olvera, M. A.; Clark, A. F.; Stone, E. M. Fibulin-5 distribution in human eyes: relevance to age-related macular degeneration. Experimental eye research 2007, 84 (2), 378–380. DOI: S0014-4835(06)00392-7 [pii] 10.1016/j.exer.2006.09.021.

(20) Vierkotten, S.; Muether, P. S.; Fauser, S. Overexpression of HTRA1 leads to ultrastructural changes in the elastic layer of Bruch’s membrane via cleavage of extracellular matrix components. PLoS One 2011, 6 (8), e22959. DOI: 10.1371/journal.pone.0022959 From NLM Medline.

(21) Schultz, D. W.; Klein, M. L.; Humpert, A. J.; Luzier, C. W.; Persun, V.; Schain, M.; Mahan, A.; Runckel, C.; Cassera, M.; Vittal, V.; et al. Analysis of the ARMD1 locus: evidence that a mutation in HEMICENTIN-1 is associated with age-related macular degeneration in a large family. Human molecular genetics 2003, 12 (24), 3315–3323. DOI: 10.1093/hmg/ddg348 From NLM Medline.

(22) Tom, I.; Pham, V. C.; Katschke, K. J., Jr.; Li, W.; Liang, W. C.; Gutierrez, J.; Ah Young, A.; Figueroa, I.; Eshghi, S. T.; Lee, C. V.; et al. Development of a therapeutic anti-HtrA1 antibody and the identification of DKK3 as a pharmacodynamic biomarker in geographic atrophy. Proceedings of the National Academy of Sciences of the United States of America 2020, 117 (18), 9952–9963. DOI: 10.1073/pnas.1917608117.

(23) Gerhardy, S.; Ultsch, M.; Tang, W.; Green, E.; Holden, J. K.; Li, W.; Estevez, A.; Arthur, C.; Tom, I.; Rohou, A.; Kirchhofer, D. Allosteric inhibition of HTRA1 activity by a conformational lock mechanism to treat age-related macular degeneration. Nature communications 2022, 13 (1), 5222. DOI: 10.1038/s41467-022-32760-9.

(24) Li, Y.; Wei, Y.; Ultsch, M.; Li, W.; Tang, W.; Tombling, B.; Gao, X.; Dimitrova, Y.; Gampe, C.; Fuhrmann, J.; et al. Cystine-knot peptide inhibitors of HTRA1 bind to a cryptic pocket within the active site region. Nature communications 2024, 15 (1), 4359. DOI: 10.1038/s41467-024-48655-w From NLM Medline.

(25) Dennis, D. G.; Joo Sun, Y.; Parsons, D. E.; Mahajan, V. B.; Smith, M. Identification of highly potent and selective HTRA1 inhibitors. Bioorg Med Chem Lett 2024, 109, 129814. DOI: 10.1016/j.bmcl.2024.129814 From NLM Medline.

(26) Song, D.; Lee, J. Y.; Park, E. C.; Choi, N. E.; Nam, H. Y.; Seo, J.; Lee, J. Structure-activity relationship analysis of activity-based probes targeting HTRA family of serine proteases. Bioorg Med Chem Lett 2023, 87, 129259. DOI: 10.1016/j.bmcl.2023.129259 From NLM Medline.

(27) Doudna, J. A.; Charpentier, E. Genome editing. The new frontier of genome engineering with CRISPR-Cas9. Science (New York, N.Y 2014, 346 (6213), 1258096. DOI: 10.1126/science.1258096.

(28) Sharma, A.; Toepfer, C. N.; Ward, T.; Wasson, L.; Agarwal, R.; Conner, D. A.; Hu, J. H.; Seidman, C. E. CRISPR/Cas9-Mediated Fluorescent Tagging of Endogenous Proteins in Human Pluripotent Stem Cells. Curr Protoc Hum Genet 2018, 96, 21 11 21-21 11 20. DOI: 10.1002/cphg.52.

(29) Leonetti, M. D.; Sekine, S.; Kamiyama, D.; Weissman, J. S.; Huang, B. A scalable strategy for high-throughput GFP tagging of endogenous human proteins. Proceedings of the National Academy of Sciences of the United States of America 2016, 113 (25), E3501–3508. DOI: 10.1073/pnas.1606731113.

(30) Schwinn, M. K.; Machleidt, T.; Zimmerman, K.; Eggers, C. T.; Dixon, A. S.; Hurst, R.; Hall, M. P.; Encell, L. P.; Binkowski, B. F.; Wood, K. V. CRISPR-Mediated Tagging of Endogenous Proteins with a Luminescent Peptide. ACS Chem Biol 2018, 13 (2), 467–474. DOI: 10.1021/acschembio.7b00549.

(31) Cho, N. H.; Cheveralls, K. C.; Brunner, A. D.; Kim, K.; Michaelis, A. C.; Raghavan, P.; Kobayashi, H.; Savy, L.; Li, J. Y.; Canaj, H.; et al. OpenCell: Endogenous tagging for the cartography of human cellular organization. Science (New York, N.Y 2022, 375 (6585), eabi6983. DOI: 10.1126/science.abi6983.

(32) Schwinn, M. K.; Steffen, L. S.; Zimmerman, K.; Wood, K. V.; Machleidt, T. A Simple and Scalable Strategy for Analysis of Endogenous Protein Dynamics. Scientific reports 2020, 10 (1), 8953. DOI: 10.1038/s41598-020-65832-1.

(33) Dixon, A. S.; Schwinn, M. K.; Hall, M. P.; Zimmerman, K.; Otto, P.; Lubben, T. H.; Butler, B. L.; Binkowski, B. F.; Machleidt, T.; Kirkland, T. A.; et al. NanoLuc Complementation Reporter Optimized for Accurate Measurement of Protein Interactions in Cells. ACS Chem Biol 2016, 11 (2), 400–408. DOI: 10.1021/acschembio.5b00753.

(34) Hageman, G. S.; Anderson, D. H.; Johnson, L. V.; Hancox, L. S.; Taiber, A. J.; Hardisty, L. I.; Hageman, J. L.; Stockman, H. A.; Borchardt, J. D.; Gehrs, K. M.; et al. A common haplotype in the complement regulatory gene factor H (HF1/CFH) predisposes individuals to age-related macular degeneration. Proceedings of the National Academy of Sciences of the United States of America 2005, 102 (20), 7227–7232.

(35) Haines, J. L.; Hauser, M. A.; Schmidt, S.; Scott, W. K.; Olson, L. M.; Gallins, P.; Spencer, K. L.; Kwan, S. Y.; Noureddine, M.; Gilbert, J. R.; et al. Complement factor H variant increases the risk of age-related macular degeneration. Science (New York, N.Y 2005, 308 (5720), 419–421. DOI: 10.1126/science.1110359.

(36) Klein, R. J.; Zeiss, C.; Chew, E. Y.; Tsai, J. Y.; Sackler, R. S.; Haynes, C.; Henning, A. K.; SanGiovanni, J. P.; Mane, S. M.; Mayne, S. T.; et al. Complement factor H polymorphism in age-related macular degeneration. Science (New York, N.Y 2005, 308 (5720), 385–389. DOI: 1109557 [pii] 10.1126/science.1109557.

(37) Despriet, D. D.; Klaver, C. C.; Witteman, J. C.; Bergen, A. A.; Kardys, I.; de Maat, M. P.; Boekhoorn, S. S.; Vingerling, J. R.; Hofman, A.; Oostra, B. A.; et al. Complement factor H polymorphism, complement activators, and risk of age-related macular degeneration. Jama 2006, 296 (3), 301–309.

(38) DiCesare, S. M.; Ortega, A. J.; Collier, G. E.; Daniel, S.; Thompson, K. N.; McCoy, M. K.; Posner, B. A.; Hulleman, J. D. GSK3 inhibition reduces ECM production and prevents age-related macular degeneration-like pathology. JCI Insight 2024, 9 (15). DOI: 10.1172/jci.insight.178050 From NLM Medline.

(39) Lankford, K. P.; Hulleman, J. D. Protocol for HiBiT tagging endogenous proteins using CRISPR-Cas9 gene editing. STAR Protoc 2024, 5 (2), 103000. DOI: 10.1016/j.xpro.2024.103000 From NLM Medline.

(40) Hazim, R. A.; Volland, S.; Yen, A.; Burgess, B. L.; Williams, D. S. Rapid differentiation of the human RPE cell line, ARPE-19, induced by nicotinamide. Experimental eye research 2019, 179, 18–24. DOI: 10.1016/j.exer.2018.10.009.

(41) Nguyen, A.; Hulleman, J. D. Evidence of Alternative Cystatin C Signal Sequence Cleavage Which Is Influenced by the A25T Polymorphism. PLoS One 2016, 11 (2), e0147684. DOI: 10.1371/journal.pone.0147684.

(42) Corso, G.; Deng, A.; Fry, B.; Polizzi, N.; Barzilay, R.; Jaakkola, T. Deep Confident Steps to New Pockets: Strategies for Docking Generalization. ArXiv 2024. From NLM PubMed-not-MEDLINE.

(43) Riching, K. M.; Mahan, S.; Corona, C. R.; McDougall, M.; Vasta, J. D.; Robers, M. B.; Urh, M.; Daniels, D. L. Quantitative Live-Cell Kinetic Degradation and Mechanistic Profiling of PROTAC Mode of Action. ACS Chem Biol 2018, 13 (9), 2758–2770. DOI: 10.1021/acschembio.8b00692 From NLM Medline.

(44) Caine, E. A.; Mahan, S. D.; Johnson, R. L.; Nieman, A. N.; Lam, N.; Warren, C. R.; Riching, K. M.; Urh, M.; Daniels, D. L. Targeted Protein Degradation Phenotypic Studies Using HaloTag CRISPR/Cas9 Endogenous Tagging Coupled with HaloPROTAC3. Curr Protoc Pharmacol 2020, 91 (1), e81. DOI: 10.1002/cpph.81.

(45) Zhang, J. H.; Chung, T. D.; Oldenburg, K. R. A Simple Statistical Parameter for Use in Evaluation and Validation of High Throughput Screening Assays. J Biomol Screen 1999, 4 (2), 67–73.

(46) Benkhaya, S.; M’Rabet, S.; El Harfi, A. Classifications, properties, recent synthesis and applications of azo dyes. Heliyon 2020, 6 (1), e03271. DOI: 10.1016/j.heliyon.2020.e03271 From NLM PubMed-not-MEDLINE.

(47) Yakupova, E. I.; Bobyleva, L. G.; Vikhlyantsev, I. M.; Bobylev, A. G. Congo Red and amyloids: history and relationship. Biosci Rep 2019, 39 (1). DOI: 10.1042/BSR20181415 From NLM Medline.

(48) He, Z.; Yan, L.; Yong, Z.; Dong, Z.; Dong, H.; Gong, Z. Chicago sky blue 6B, a vesicular glutamate transporters inhibitor, attenuates methamphetamine-induced hyperactivity and behavioral sensitization in mice. Behav Brain Res 2013, 239, 172–176. DOI: 10.1016/j.bbr.2012.11.003 From NLM Medline.

(49) Peng, H.; Hulleman, J. D. Prospective Application of Activity-Based Proteomic Profiling in Vision Research-Potential Unique Insights into Ocular Protease Biology and Pathology. International journal of molecular sciences 2019, 20 (16). DOI: 10.3390/ijms20163855.

(50) Cravatt, B. F.; Wright, A. T.; Kozarich, J. W. Activity-based protein profiling: from enzyme chemistry to proteomic chemistry. Annu Rev Biochem 2008, 77, 383–414. DOI: 10.1146/annurev.biochem.75.101304.124125.

(51) Liu, Y.; Patricelli, M. P.; Cravatt, B. F. Activity-based protein profiling: the serine hydrolases. Proceedings of the National Academy of Sciences of the United States of America 1999, 96 (26), 14694–14699.

(52) Bachovchin, D. A.; Cravatt, B. F. The pharmacological landscape and therapeutic potential of serine hydrolases. Nat Rev Drug Discov 2012, 11 (1), 52–68. DOI: 10.1038/nrd3620.

(53) Corso, G., Stärk, H., Jing, B., Barzilay, R., Jaakkola, T. DiffDock: Diffusion Steps, Twists, and Turns for Molecular Docking. arXiv 2022. DOI: 10.48550/arXiv.2210.01776.

(54) Joyner, A. L.; Sabin, F. R. Altered Cutaneous Conditions in the Skin of Tuberculous Guinea Pigs as Demonstrated with a Vital Dye. The Journal of experimental medicine 1938, 68 (3), 325–334. DOI: 10.1084/jem.68.3.325 From NLM PubMed-not-MEDLINE.

(55) Weinberg, J.; Greaney, E. M.; Rawlings, B.; Haley, T. J. The use and toxicity of Pontamine Sky Blue. Science (New York, N.Y 1951, 114 (2950), 41–42. DOI: 10.1126/science.114.2950.41.

(56) Weinberg, J.; Greaney, E. M. Identification of regional lymph nodes by means of a vital staining dye during surgery of gastric cancer. Surg Gynecol Obstet 1950, 90 (5), 561–567. From NLM Medline.

(57) Yifa, O.; Weisinger, K.; Bassat, E.; Li, H.; Kain, D.; Barr, H.; Kozer, N.; Genzelinakh, A.; Rajchman, D.; Eigler, T.; et al. The small molecule Chicago Sky Blue promotes heart repair following myocardial infarction in mice. JCI Insight 2019, 4 (22). DOI: 10.1172/jci.insight.128025 From NLM Medline.

(58) Min, J. O.; Strohaker, T.; Jeong, B. C.; Zweckstetter, M.; Lee, S. J. Chicago sky blue 6B inhibits alpha-synuclein aggregation and propagation. Mol Brain 2022, 15 (1), 27. DOI: 10.1186/s13041-022-00913-y From NLM Medline.

(59) Pomierny, B.; Krzyzanowska, W.; Skorkowska, A.; Jurczyk, J.; Bystrowska, B.; Budziszewska, B.; Pera, J. Inhibition of Vesicular Glutamate Transporters (VGLUTs) with Chicago Sky Blue 6B Before Focal Cerebral Ischemia Offers Neuroprotection. Mol Neurobiol 2023, 60 (6), 3130–3146. DOI: 10.1007/s12035-023-03259-1 From NLM Medline.

(60) Smeralda, W.; Since, M.; Corvaisier, S.; Fayolle, D.; Cardin, J.; Duprey, S.; Jourdan, J. P.; Cullin, C.; Malzert-Freon, A. A Biomimetic Multiparametric Assay to Characterise Anti-Amyloid Drugs. International journal of molecular sciences 2023, 24 (23). DOI: 10.3390/ijms242316982 From NLM Medline.

(61) Pomierny, B.; Krzyzanowska, W.; Skorkowska, A.; Jurczyk, J.; Budziszewska, B.; Pera, J. Chicago sky blue 6B exerts neuroprotective and anti-inflammatory effects on focal cerebral ischemia. Biomedicine & pharmacotherapy = Biomedecine & pharmacotherapie 2024, 170, 116102. DOI: 10.1016/j.biopha.2023.116102 From NLM Medline.

(62) Chen, S.; Puri, A.; Bell, B.; Fritsche, J.; Palacios, H. H.; Balch, M.; Sprunger, M. L.; Howard, M. K.; Ryan, J. J.; Haines, J. N.; et al. HTRA1 disaggregates alpha-synuclein amyloid fibrils and converts them into non-toxic and seeding incompetent species. Nature communications 2024, 15 (1), 2436. DOI: 10.1038/s41467-024-46538-8 From NLM Medline.

(63) Munoz, S. S.; Li, H.; Ruberu, K.; Chu, Q.; Saghatelian, A.; Ooi, L.; Garner, B. The serine protease HtrA1 contributes to the formation of an extracellular 25-kDa apolipoprotein E fragment that stimulates neuritogenesis. The Journal of biological chemistry 2018, 293 (11), 4071–4084. DOI: 10.1074/jbc.RA117.001278 From NLM Medline.

(64) Fukutake, T. Cerebral autosomal recessive arteriopathy with subcortical infarcts and leukoencephalopathy (CARASIL): from discovery to gene identification. J Stroke Cerebrovasc Dis 2011, 20 (2), 85–93. DOI: 10.1016/j.jstrokecerebrovasdis.2010.11.008 From NLM Medline.

(65) Uemura, M.; Nozaki, H.; Koyama, A.; Sakai, N.; Ando, S.; Kanazawa, M.; Kato, T.; Onodera, O. HTRA1 Mutations Identified in Symptomatic Carriers Have the Property of Interfering the Trimer-Dependent Activation Cascade. Front Neurol 2019, 10, 693. DOI: 10.3389/fneur.2019.00693 From NLM PubMed-not-MEDLINE.

(66) Al-Tohamy, R.; Ali, S. S.; Li, F.; Okasha, K. M.; Mahmoud, Y. A.; Elsamahy, T.; Jiao, H.; Fu, Y.; Sun, J. A critical review on the treatment of dye-containing wastewater: Ecotoxicological and health concerns of textile dyes and possible remediation approaches for environmental safety. Ecotoxicol Environ Saf 2022, 231, 113160. DOI: 10.1016/j.ecoenv.2021.113160 From NLM Medline.

(67) Misasi, R.; Huemer, H. P.; Schwaeble, W.; Solder, E.; Larcher, C.; Dierich, M. P. Human complement factor H: an additional gene product of 43 kDa isolated from human plasma shows cofactor activity for the cleavage of the third component of complement. European journal of immunology 1989, 19 (9), 1765–1768. DOI: 10.1002/eji.1830190936.

(1) Liu, J. Y.; Zhu, Y. C.; Zhou, L. X.; Wei, Y. P.; Mao, C. H.; Cui, L. Y.; Peng, B.; Yao, M. HTRA1-related autosomal dominant cerebral small vessel disease. Chin Med J (Engl) 2020, 134 (2), 178–184. DOI: 10.1097/CM9.0000000000001176 From NLM Medline.

(2) Gerhardy, S.; Ultsch, M.; Tang, W.; Green, E.; Holden, J. K.; Li, W.; Estevez, A.; Arthur, C.; Tom, I.; Rohou, A.; Kirchhofer, D. Allosteric inhibition of HTRA1 activity by a conformational lock mechanism to treat age-related macular degeneration. Nature communications 2022, 13 (1), 5222. DOI: 10.1038/s41467-022-32760-9 From NLM Medline.

(3) Nozaki, H.; Kato, T.; Nihonmatsu, M.; Saito, Y.; Mizuta, I.; Noda, T.; Koike, R.; Miyazaki, K.; Kaito, M.; Ito, S.; et al. Distinct molecular mechanisms of HTRA1 mutants in manifesting heterozygotes with CARASIL. Neurology 2016, 86 (21), 1964–1974. DOI: 10.1212/WNL.0000000000002694 From NLM Medline.

(4) Uemura, M.; Nozaki, H.; Koyama, A.; Sakai, N.; Ando, S.; Kanazawa, M.; Kato, T.; Onodera, O. HTRA1 Mutations Identified in Symptomatic Carriers Have the Property of Interfering the Trimer-Dependent Activation Cascade. Front Neurol 2019, 10, 693. DOI: 10.3389/fneur.2019.00693 From NLM PubMed-not-MEDLINE.

