## Supplemental Information for "Select azo compounds post-translationally modulate HTRA1 abundance and activity potentially through interactions at the trimer interface"

**Supplemental Figure 1.** Sanger sequence alignment of the HTRA1 exon 9 edited (top) band shown in Fig. 1B. The full VS HiBiT GG HA sequence was fully accounted for.

**Supplemental Figure 2.** Design and validation of a HiBiT CFH ARPE-19 cell line. (A,B) A 2xFLAG VS HiBiT sequence immediately after the predicted signal sequence cleavage site (determined by SignalP software, <https://services.healthtech.dtu.dk/service.php?SignalP-5.0>). The 2xFLAG VS HiBiT tag had no predicted effect on signal peptide cleavage. (C) CFH editing design. (D) CFH exon 1 gDNA amplification of unedited and edited ARPE-19 cells. Insertion of the 2xFLAG VS HiBiT sequence increases the predicted molecular weight by 87 bp. Estimated editing efficiency (top band/bottom band) is ~55%. (E) Validation of HiBiT insertion at the CFH locus by siRNA. ARPE-19 HiBiT CFH edited cells were reverse transfected with control siRNAs (non-targeting, siTOX), or siRNAs against CFH. n = 3 independent experiments. **** p < 0.0001, one-way ANOVA with multiple comparisons vs. non-targeting. (F) HiBiT tagging the CFH locus produces a full length, ~150 kDa isoform, consisting of 20 complement control protein (CCP) domains, and a smaller isoform (~50 kDa, factor H like protein 1 (FHL-1)) which contains the first 7 CCP domains of CFH^67^. (G) Nic differentiation does not alter CFH secretion at early timepoints. ARPE-19 HiBiT CFH cells were plated at confluence followed by incubation in either full DMEM/F12 or Nic media for up to 7 days. A HiBiT assay was performed on media aliquots for 1 week. n = 3 independent experiments. **** p < 0.0001, multiple unpaired t-test vs. respective DMEM/F12 control.

**Supplemental Figure 3.** Sanger sequence alignment of the CFH exon 1 edited (top) band shown in Sup. Fig. 2D. The full 2xFLAG VS HiBiT sequence was fully accounted for.

**Supplemental Figure 4.** (A-E) Chemical structures of mono azo dyes and VGLUT1 inhibitor, bromocriptine mesylate. (F) ARPE-19 HTRA1 HiBiT cells grown in DMEM/F12 full media were treated with the indicated compound (20 μM) for 72 h followed by a HiBiT assay. Representative data of 3 independent experiments.

**Supplemental Figure 5.** Full images of all gels and blots used throughout Figs. 1-5 and Sup. Fig. 2.

**Supplemental Figure 6.** (A) DiffDock confidence scores for the 10 attempted models for Chicago Sky Blue (CSB), Congo Red (Congo R), and Brilliant Yellow (Bril Y). A lower score is indicative of higher confidence. (B) Structural representations of the identified binding interfaces by DiffDock for CSB, CongoR, and Bril Y, respectively.
