## Supplemental Figures for "Select azo compounds post-translationally modulate HTRA1 abundance and activity potentially through interactions at the trimer interface"

Sup. Fig. 1

HTRA1 HiBiT

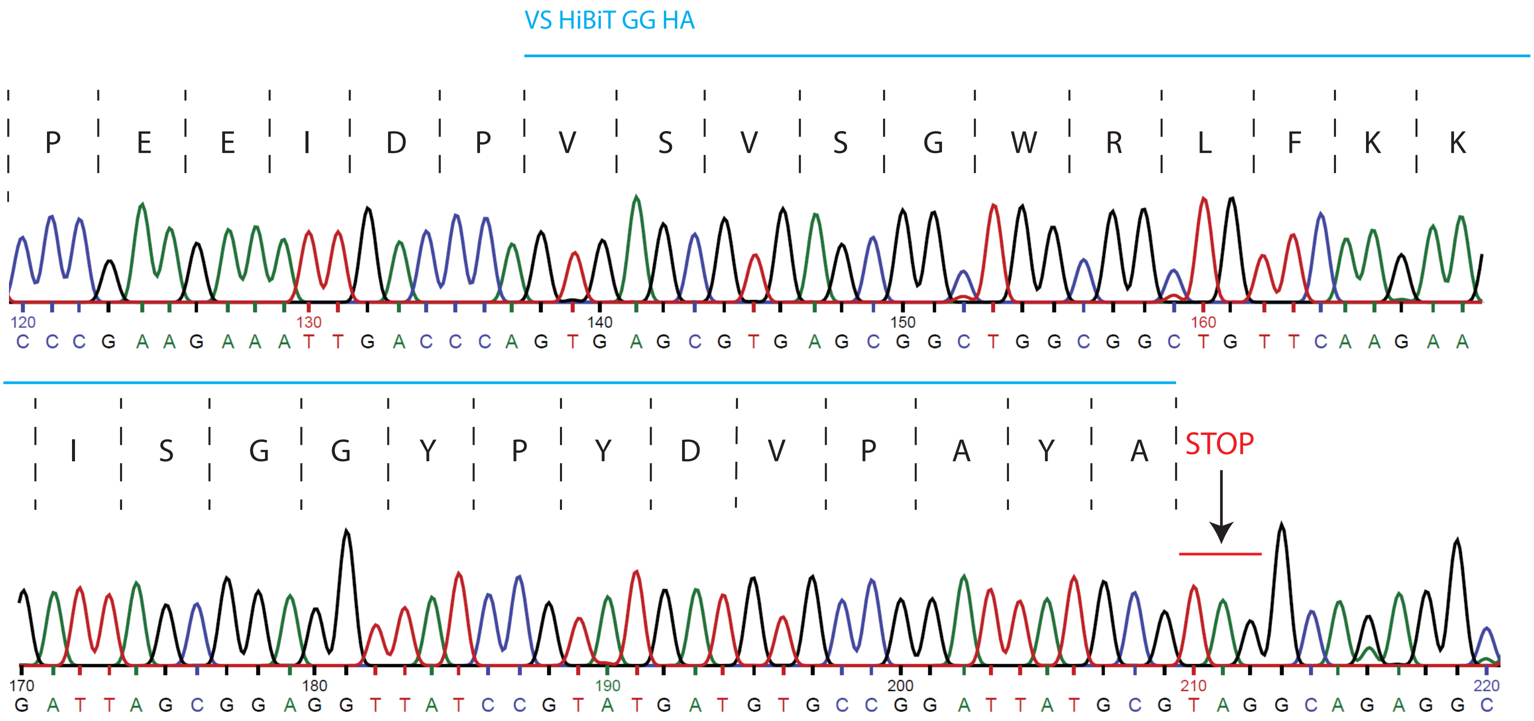

Sup. Fig. 2

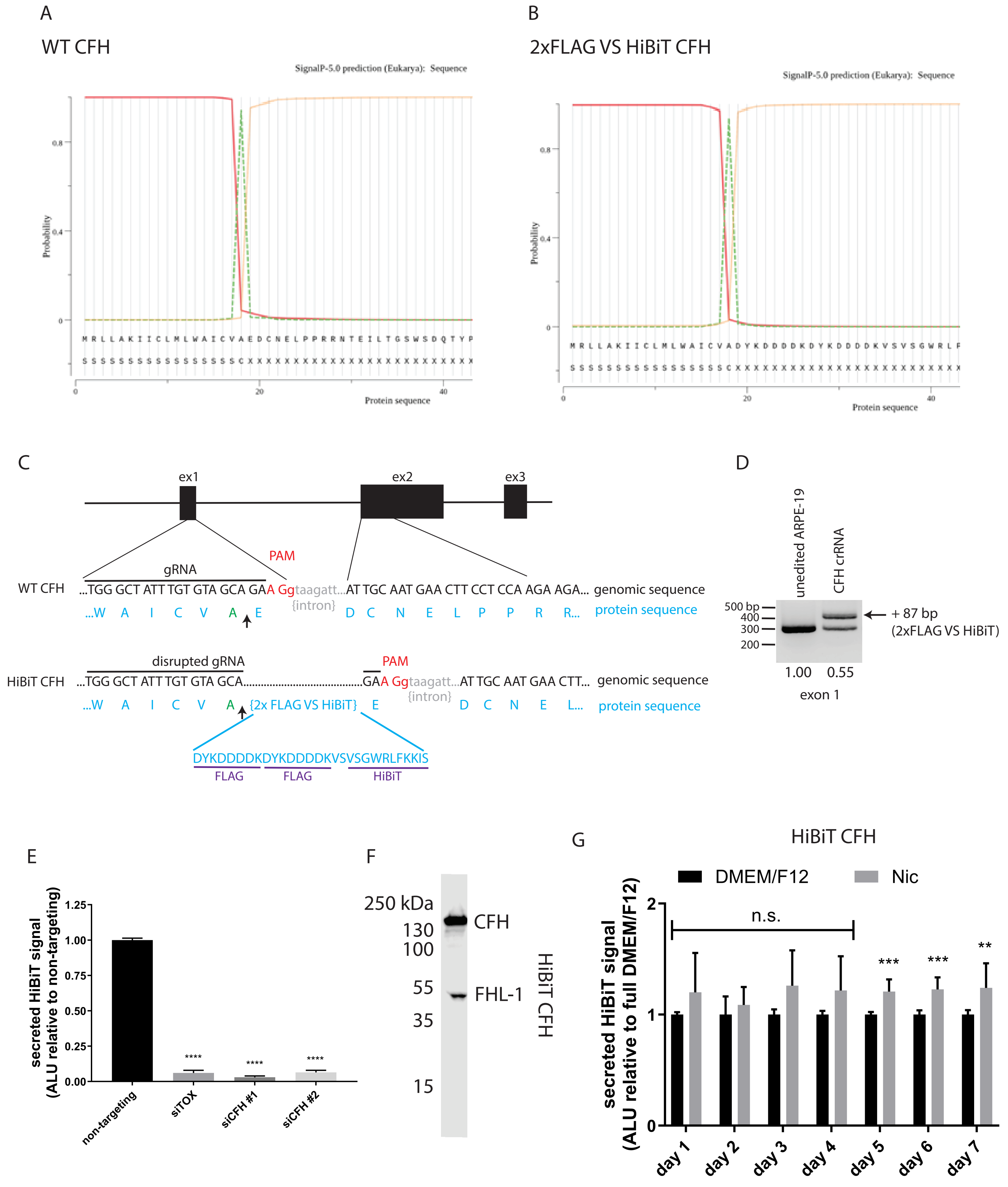
A
WT CFH
SignalP-5.0 prediction (Eukarya): Sequence
Probability
Protein sequence
M R L L A K I I C L M L W A I C V A E D C N E L P P R R N T E I L T G S W S D Q T Y P
B
2xFLAG VS HiBiT CFH
SignalP-5.0 prediction (Eukarya): Sequence
M R L L A K I I C L M L W A I C V A D Y K D D D D K D Y K D D D D K V S V S G W R L F
C
ex1
ex2
ex3
gRNA
PAM
WT CFH
...TGG GCT ATT TGT GTA GCA GAA Ggtaagatt...AT TGC AAT GAA CTT CCT CCA AGA AGA...
genomic sequence
{intron}
...W A I C V A E D C N E L P P R R...
protein sequence
disrupted gRNA
HiBiT CFH
...TGG GCT ATT TGT GTA GCA...GAA Ggtaagatt...AT TGC AAT GAA CTT...
...W A I C V A {2x FLAG VS HiBiT} E D C N E L...
DYKDDDDKDYKDDDDKVSVSGWRLFKKIS
FLAG
FLAG
HiBiT
D
unedited ARPE-19
CFH crRNA
500 bp
400
300
200
+ 87 bp
(2xFLAG VS HiBiT)
1.00
0.55
exon 1
E
secreted HiBiT signal
(ALU relative to non-targeting)
1.25
1.00
0.75
0.50
0.25
0.00
****
non-targeting
siTOX
siCFH #1
siCFH #2
F
250 kDa
130
100
55
35
15
CFH
FHL-1
HiBiT CFH
G
HiBiT CFH
DMEM/F12
Nic
n.s.
***
**
secreted HiBiT signal
(ALU relative to full DMEM/F12)
day 1
day 2
day 3
day 4
day 5
day 6
day 7

Sup. Fig. 3

HiBiT CFH

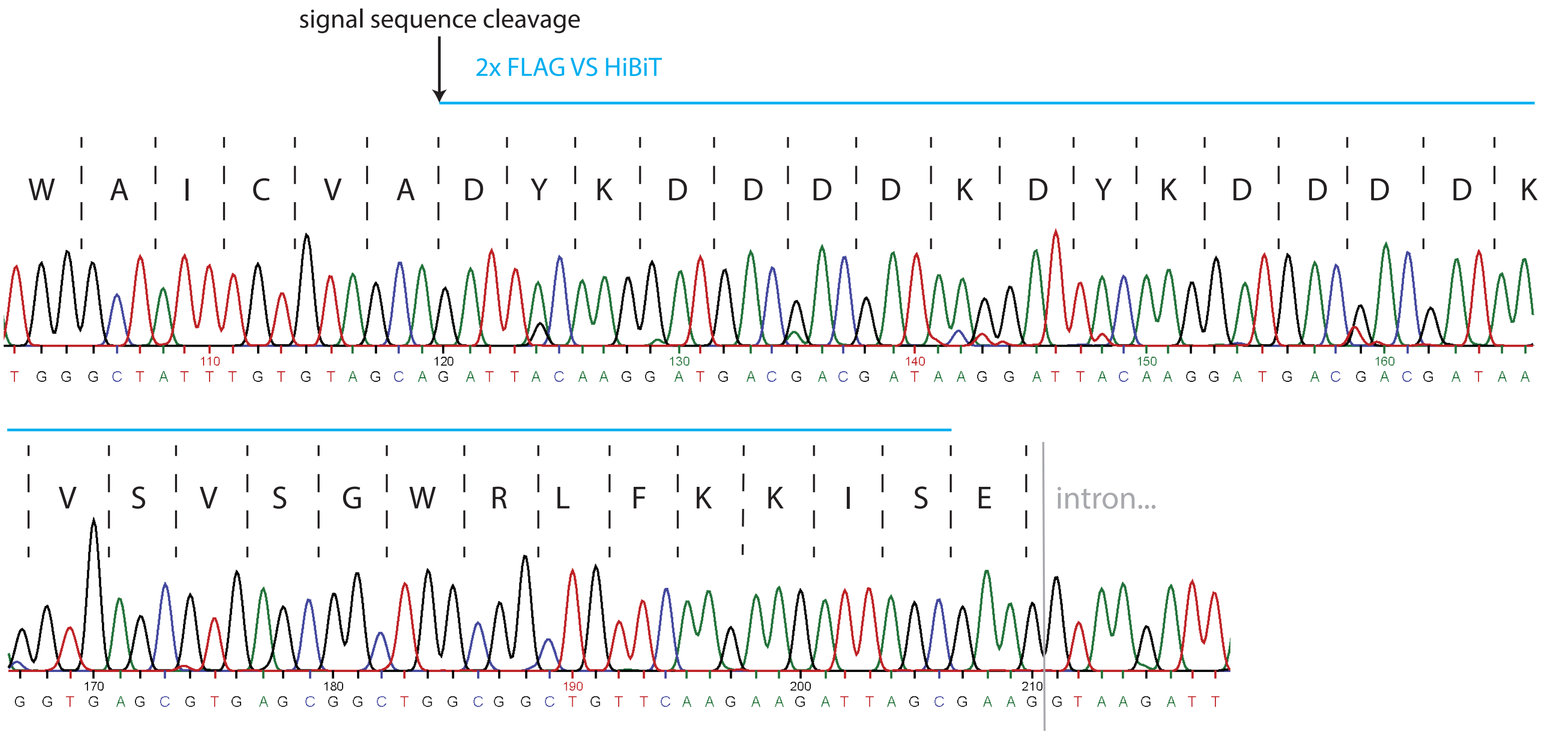

Sup. Fig. 4

A

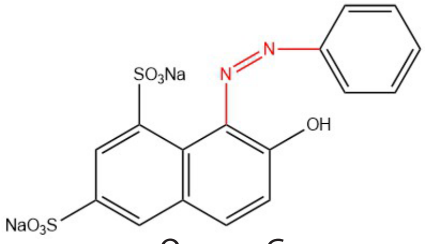


Orange G

B

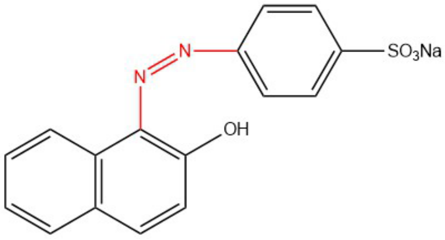


Orange 2

C

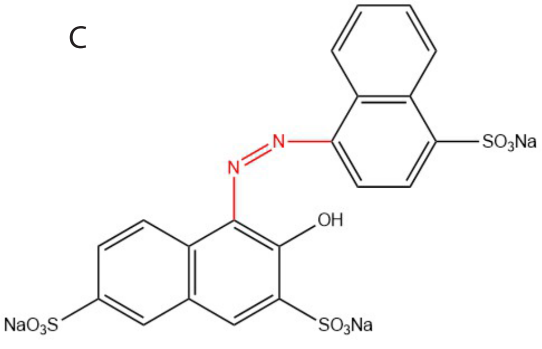


Amaranth

D

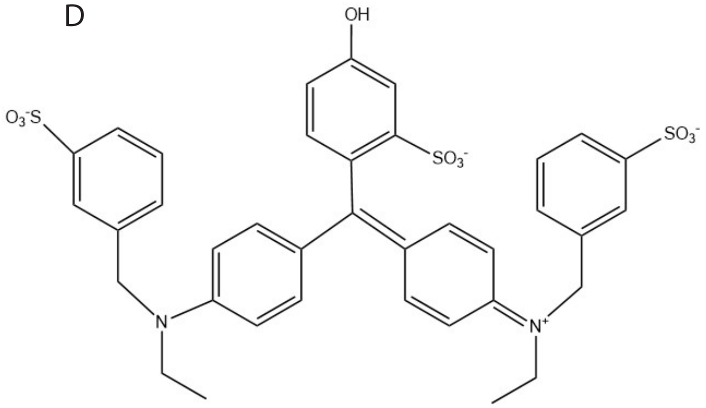


Fast Green (control dye)

E

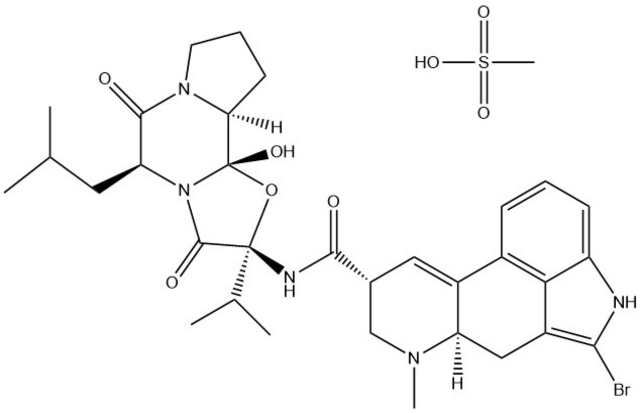


Bromocriptine mesylate (VGLUT1 inhibitor)

F

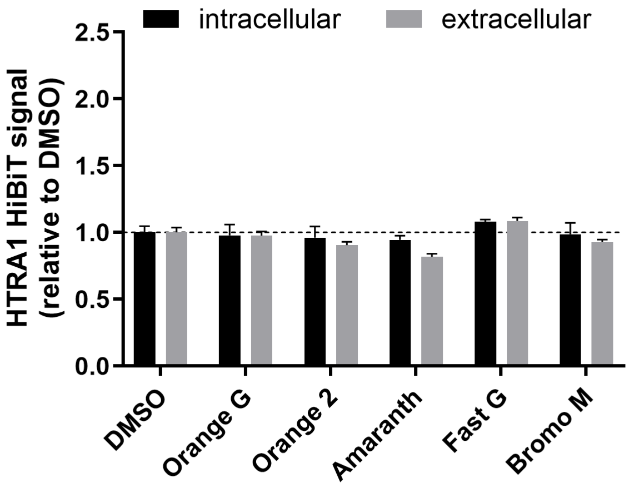

Sup. Fig. 5

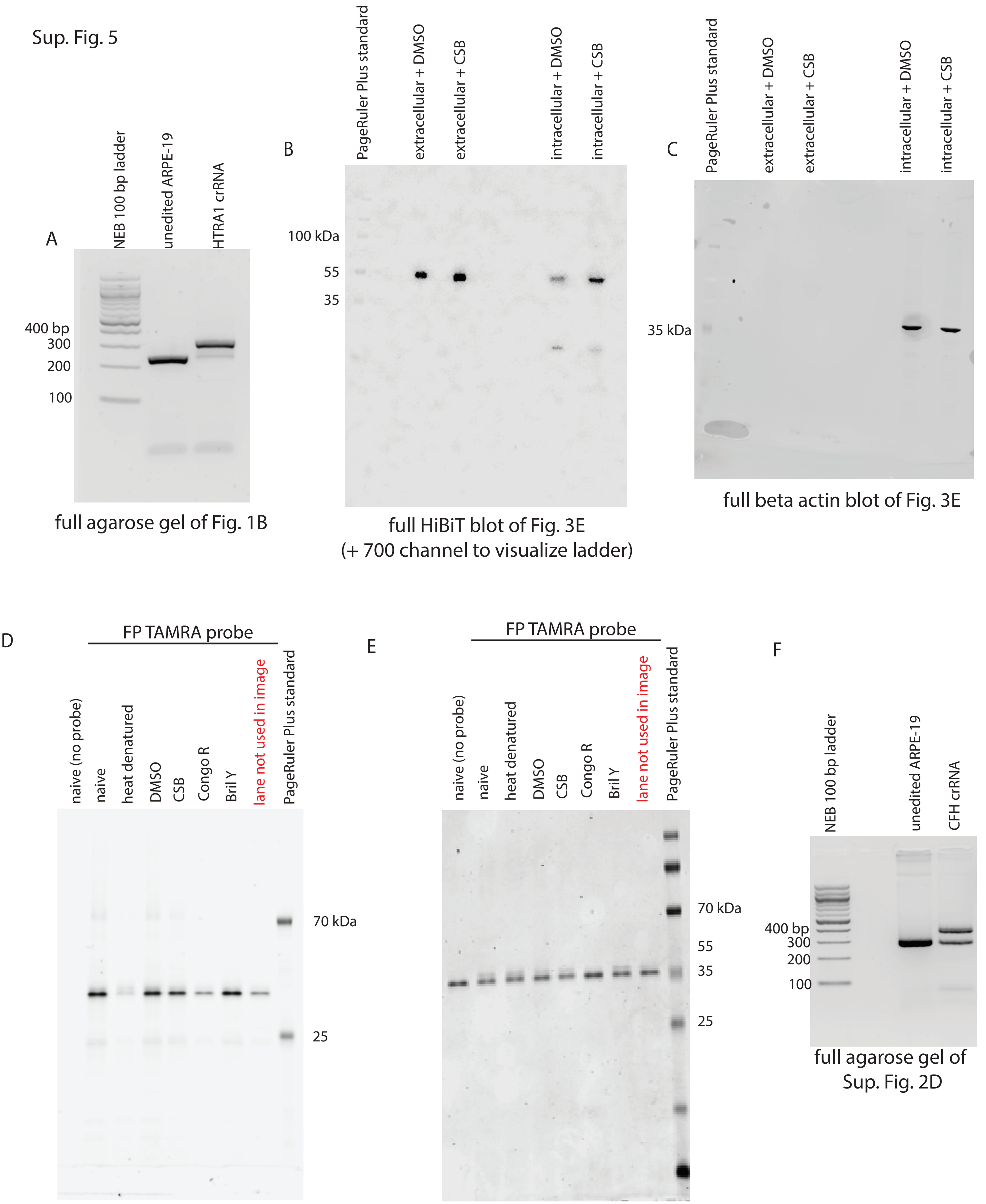


full agarose gel of Fig. 1B

full HiBiT blot of Fig. 3E
(+ 700 channel to visualize ladder)

full beta actin blot of Fig. 3E

full SDS-PAGE gel (TAMRA channel)
of Fig. 5A

full SDS-PAGE gel (Coomassie channel)
of Fig. 5A

full agarose gel of
Sup. Fig. 2D

Sup. Fig. 6

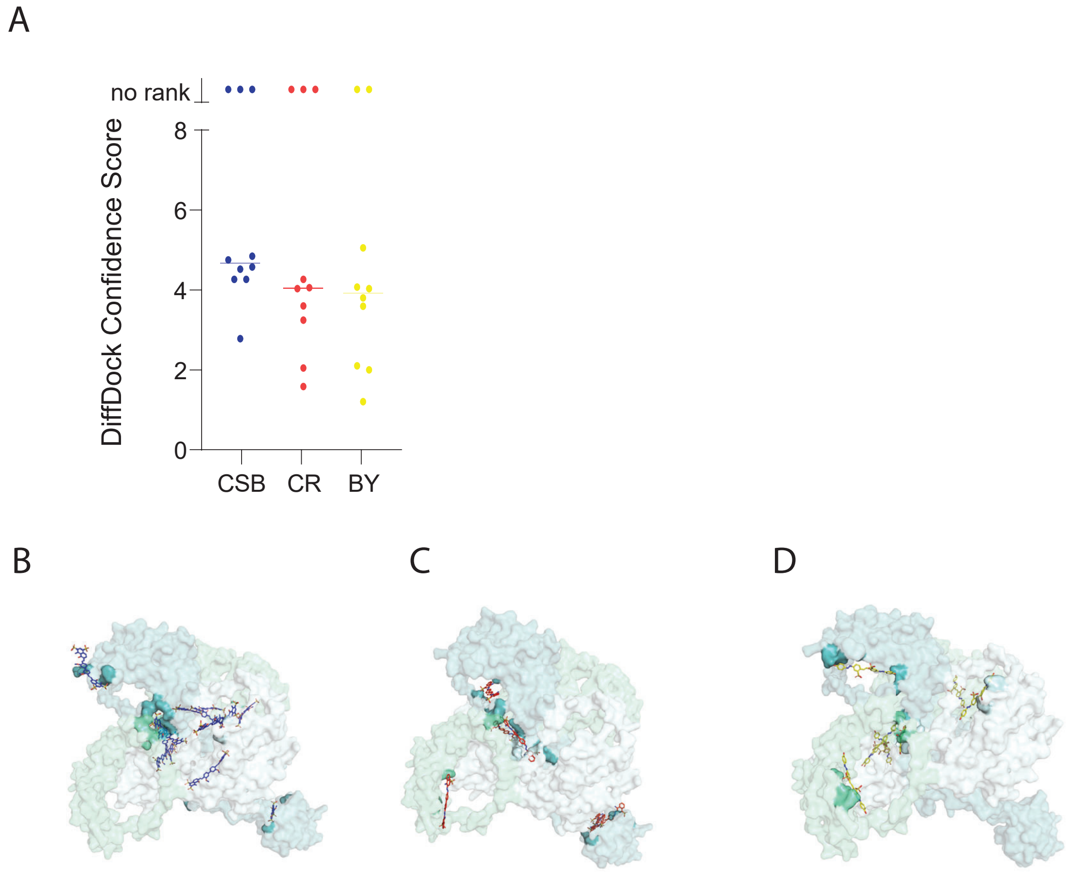

Table S1:

| HiBiT edit | gRNA | PAM | ssODN |
|---|---|---|---|
| VS HiBiT GG HTRA1 | TCCCGAAGAAATTGACCCAT | AGG | GGGGTAATGAAGATATCATGATCACAGTGATTCCCGAAGAAATTGACCCAGTGAGCGTGAGCGGCTGGCGGCTGTTCAAGAAGATTAGCGGAGGTTATCCGTATGATGTGCCGGATTATGCGTAGGCAGAGGCATGAGCTGGACTTCATGTTTCCCTCAAAGACTCTCCCGT |
| 2xFLAG VS HiBiT CFH | TGGGCTATTTGTGTAGCAGA | AGG | ATGAGACTTCTAGCAAAGATTATTTGCCTTATGTTATGGGCTATTTGTGTAGCAGATTACAAGGATGACGACGATAAGGATTACAAGGATGACGACGATAAGGTGAGCGTGAGCGGCTGGCGGCTGTTCAAGAAGATTAGCGAAGGTAAGATTAAAAGAGACTCTTTTCTGAAAACTGTATTATGAAACATTTGCT |

Table S2:

| **HiBiT edit** | **exon** | **forward primer** | **reverse primer** |
|---|---|---|---|
| VS HiBiT GG HTRA1 | 9 | GCAGTGGTGGTCTCAAGGAAAACGACGTCATAATCA | CAGAGTCCTCATCCGTCATCCACGGGAGAGTCTTT |
| 2xFLAG VS HiBiT CFH | 1 | TTGCAGCAAGTTCTTTCCTGCACTAATCACAATTCTTG | GCCACTCAATTGTCAAGTTACAGAATACTTAAGCACATT |

Table S3:

| Target | sense sequence |
|---|---|
| siTOX transfection control | proprietary |
| non-targeting siRNA control | proprietary |
| siHtrA1 #1 | AUAUCGAAUUGUUUCGCAAtt |
| siHtrA1 #2 | UGAUCACGUGAUUCCCGAtt |
| siHtrA2 | GCACCUGCCGUGGUCUAUAtt |
| siHtrA3 | CAAGAUCCAUCCCAAGAAAtt |
| siCFH #1 | GGAUAUAGAUCUCUUGGAAtt |
| siCFH #2 | CCAGAUCGGGAAUACCAUUtt |
